# Evolutionary history of Jamestown Canyon virus disentangles complex multi-vector ecology

**DOI:** 10.64898/2026.01.09.698726

**Authors:** Ellie Bourgikos, Simon Dellicour, Philippe Lemey, Nicole M. Feriancek, Angela B. Bransfield, Mallery I. Breban, Michael J. Misencik, Tanya A. Petruff, John J. Shepard, Theodore G. Andreadis, John F. Anderson, Kiet A. Ngo, Joseph G. Maffei, Alan P. Dupuis, Stephen M. Rich, Guang Xu, Gabrielle Sakolsky, Keith J. Price, Madeline L. Metzger, Marc A. Suchard, Rafael Lopes, Fabiana Gámbaro, Colin J. Carlson, Guy Baele, Alexander T. Ciota, Chantal B.F. Vogels, Verity Hill, Philip M. Armstrong, Nathan D. Grubaugh

## Abstract

Jamestown Canyon virus (JCV) is a re-emerging mosquito-borne virus of increasing concern in North America. It has been historically understudied, leading to significant gaps in our understanding of its evolutionary history, ecological maintenance, and transmission dynamics. Here, we generated 658 whole-genome JCV sequences from the Northeast United States, including 84% (500/597) of all JCV-positive mosquitoes detected in Connecticut from 1997-2022. Then we applied phylodynamic methods to demonstrate how mosquito phenology and host interaction structure the persistence and spread of JCV. Our phylogenetic analyses estimate that JCV was introduced in the Northeast by at least the early 1700s and the primary introductions of lineages A and B into Connecticut occurred during the mid-1800s to mid-1900s. Further, we estimate that JCV evolves at a rate of ∼3 x 10^-5^ s/s/y, making it one of the slowest evolving known RNA viruses, because the virus spends ∼10 months per year in evolutionary stasis while over-wintering in mosquito eggs. To investigate ecological drivers of JCV spread in Connecticut, we paired discrete trait and continuous phylogeographic reconstructions with mosquito surveillance data. We estimate that JCV has a low diffusion rate of ∼30-60 km^2^/year, which is more similar to slow-moving tick-borne viruses than other mosquito-borne viruses. We found that univoltine *Aedes* mosquitoes were likely to maintain the virus across years through overwintering in eggs, accounting for its slow evolution and dispersal, while multivoltine mosquitoes contribute to periodic bursts of spatial diffusion and amplification within seasons. By characterizing seasonal dynamics of JCV, we demonstrate the utility of dense sequencing and phylodynamics to disentangle complex transmission cycles, offering a framework to rapidly advance our evolutionary and ecological knowledge of understudied viruses.

## Introduction

Arthropod-borne viruses, or arboviruses, represent a diverse range of pathogens impacting the health of humans and animals^1–3^. Many arbovirus species remain understudied due to their complex transmission cycles (e.g., wide-range of invertebrate vectors and vertebrate hosts) that present challenges to traditional epidemiological methods^4,5^. Among these species, Jamestown Canyon virus (family: *Peribunyaviridae*, genus: *Orthobunyavirus*, California serogroup) is a re-emerging pathogen garnering increased awareness as one of the most widespread mosquito-borne viruses in North America^6–8^. Since 2011, there have been 336 diagnosed human cases of JCV in the United States (US), distributed across 26 states, with the highest burden of cases in the Midwest and Northeast^9^. These cases have resulted in 227 hospitalizations due to neuroinvasive disease and 12 deaths^9^. Studies from throughout the endemic range of North America indicate that JCV seroprevalence in humans is ∼20%^10–16^, suggesting that many infections are asymptomatic^6,7^ and cases are severely underdiagnosed^17,18^. Despite its widespread distribution and growing public health significance, JCV remains understudied, particularly in terms of its transmission dynamics and life cycle in mosquitoes. This lack of foundational knowledge limits our ability to assess human risk, guide surveillance efforts, and implement effective control strategies, challenges emblematic of the broader difficulties in studying neglected arboviruses.

Viral phylodynamics offers a framework for investigating the evolutionary and ecological processes underlying complex arbovirus transmission cycles^19^. By analyzing the genetic relationships between virus sequences, these methods can reveal how a virus moves across space and through hosts over time. The limited availability of complete JCV genome sequences has restricted study of its evolutionary dynamics in the US. This is further complicated by its seasonal transmission cycle, which has yet to be fully characterized^20^, as the virus is maintained between mosquito and white-tailed deer populations^21^. JCV has been detected in 26 different mosquito species^17,20^, across the *Aedes*, *Culex*, *Culiseta*, and *Coquillettidia* genera^7,10^, making it ecologically diffuse and difficult to define. To gain a more comprehensive understanding of evolutionary and ecological dynamics underlying JCV transmission, our study addresses the following questions through a viral phylodynamic framework and a rich sample set from the Northeast US: 1) How is JCV evolving and what is its existing diversity? 2) When was JCV introduced to the Northeast and how did it spread across landscapes? 3) Which mosquito species are critical to JCV maintenance and diffusion?

Through dense historical sequencing of JCV, we generated a unique genomic dataset that facilitates phylodynamic analysis of the virus at a fine spatial scale. In linking 658 new virus sequences, spanning the small (S), medium (M), and large (L) genome segments, to comprehensive surveillance metadata, we are able to study the evolution of JCV across temporal and ecological dimensions. Using phylodynamic and phylogeographic methods^19^, we describe the decades-long circulation of JCV in the Northeastern US, particularly in Connecticut; the slow evolution and spread of two highly divergent virus lineages demonstrates spatial heterogeneities, suggesting ecological restrictions to spread. From our analyses, we propose that single-brood (univoltine) mosquitoes are primarily responsible for long-term trans-seasonal persistence of JCV (‘maintenance cycle’), while multi-brood (multivoltine) mosquitoes drive within-season transmission and spread (‘diffusion cycle’; **Fig. 1**). Through this work, we outline the integration of a modern high-throughput sequencing and phylodynamic framework to elucidate complex ecological cycles and transmission dynamics of a historically understudied virus with emerging public health significance.

**Figure 1.**
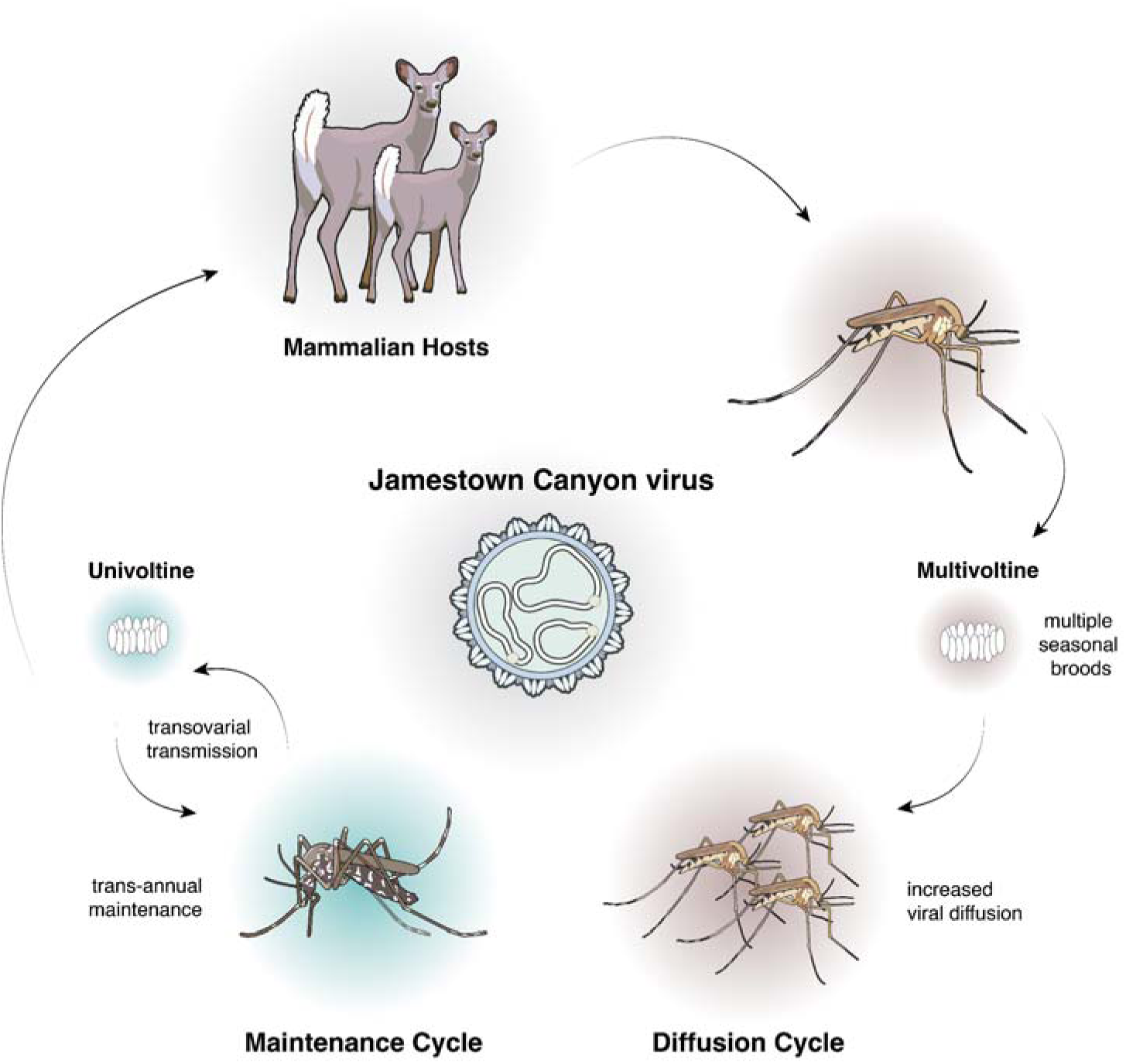
Proposed ecological cycle of Jamestown Canyon virus in the Northeast. The Maintenance Cycle (left), in blue, depicts *Aedes* mosquitoes and their eggs. As part of this cycle, univoltine *Aedes* mosquitoes occupy woodland breeding sites, with restricted spatial expansion within seasons. Synchronized spring emergence and vertical transmission facilitate long-term viral maintenance. The Diffusion Cycle (right), in brown, depicts *Anopheles* mosquitoes and their eggs. As part of this cycle multivoltine *Anopheles* mosquitoes experience extended emergence periods that enable transmission and short-term amplification within seasons. This cycle ultimately represents dead-end JCV diffusion. All images were generated using NIH BioArt.

## Results

### Genetic diversity of JCV in the United States

Historically, JCV sequencing from mosquitoes has been limited, restricting potential analysis of virus evolution and transmission. Prior to this study, only 39 whole genome sequences (i.e. sequences from all three genome segments) from JCV-positive mosquitoes were publicly available (**Fig. 2a**), of which 32 were generated by a recent study from New York^22^. While there are approximately 100 S sequences available on GenBank, predominantly from Connecticut^6,23^, whole genome sequences are required to accurately reconstruct the evolutionary history of JCV^24^, including reassortment.

**Figure 2.**
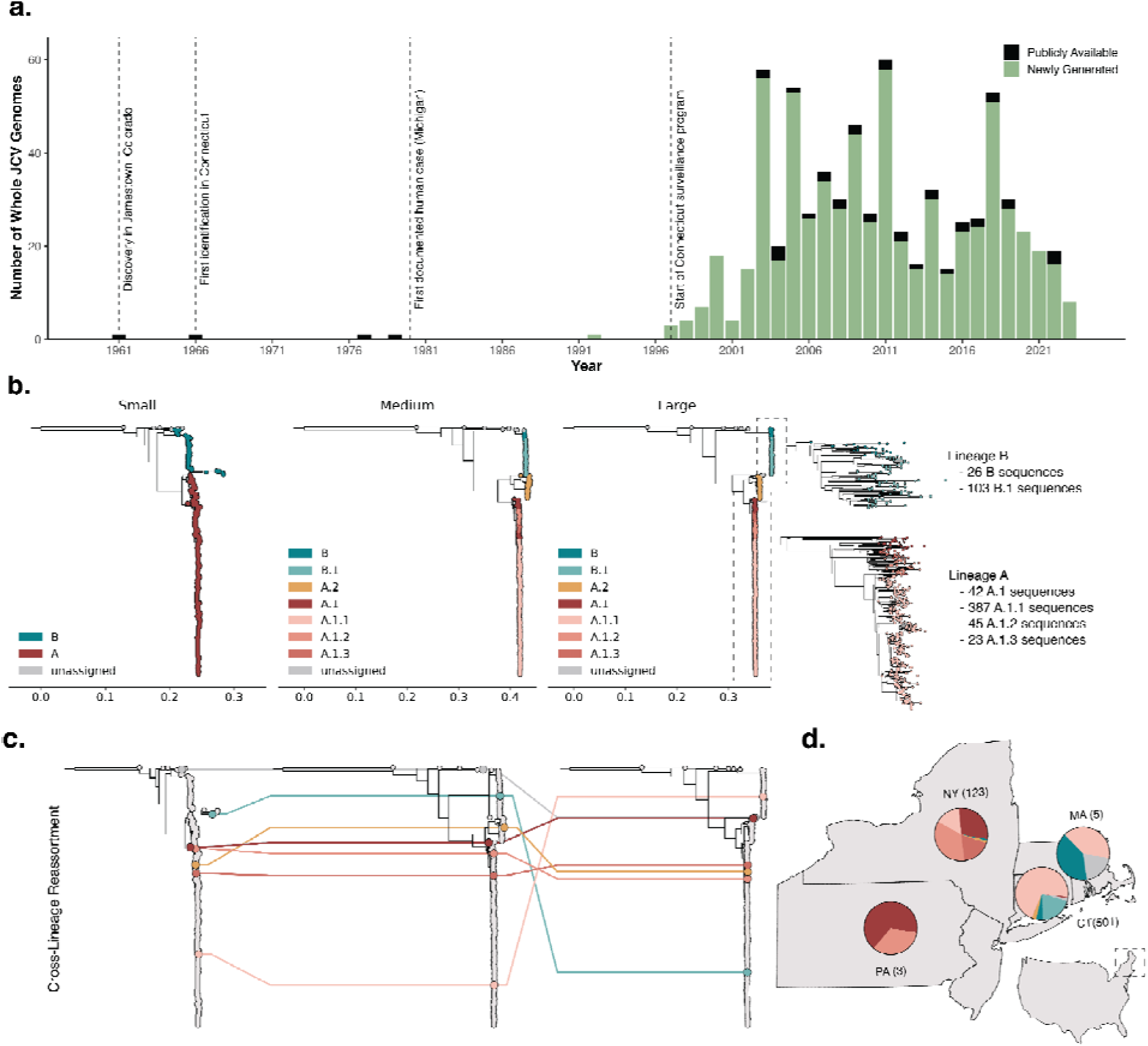
Expansion of sequencing and development of a lineage system shows JCV diversity in the Northeast US. (**a**) Availability of whole Jamestown Canyon virus (JCV) genomes from mosquitoes, defined as 70% consensus coverage for each segment; (**b**) Putative lineage system for JCV across genomic segments shown with maximum-likelihood phylogenies. Tips colored by major lineage on the S segment, and minor lineage on the M and L segments. Enlarged visualizations of the A and B clades of the L segment, colored by minor lineage, shown on right; (**c**) Tanglegram of maximum-likelihood phylogenies of the S, M, and L JCV segments. Lines connect tips of a single sampled genome across each segment, colored by the lineage of the M segment. Reassortant sequences demonstrate incongruent topology when compared across segments; (**d**) Spatial distribution of JCV lineages in the Northeast by state: Connecticut (CT) 501 sequences, New York (NY) 123 sequences, Massachusetts (MA) 5 sequences, and Pennsylvania (PA) 3 sequences.

To enhance the existing genomic dataset, we obtained 755 JCV-positive mosquito pools from 1966 to 2023 in collaboration with mosquito surveillance organizations and public health laboratories across 5 states. In processing these samples, we designed unique primers spanning all three virus segments (**Supplementary Table 1**) to be incorporated into a highly multiplexed amplicon-based sequencing method^25^, ‘JCVSeq’. Our method generated >70% of the genome from samples containing at least 193, 79, and 74 RNA copies per μL for the S, M, and L segments, respectively, and thus we were able to generate whole genomes from 87% (658/755) of the samples that we sequenced (**Extended Data Fig. 1**). We further validated our protocol using untargeted metagenomic sequencing on a smaller subset of samples, eliminating the possibility of primer mismatches in poor sequencing results (**Methods**). As a result, we generated 658 JCV sequences with >70% coverage across each of the S, M, and L genome segments (**Fig. 2a**), including sequencing the M and L segments for the partial S segments that were previously available. In total, we provide 500 novel sequences from Connecticut (1992-2022), 91 from New York (2003-2022), 60 from North Dakota (2003-2006), 4 from Massachusetts (2023), and 3 from Pennsylvania (2023).

Our expanded dataset enables us to explore and characterize JCV genetic diversity at a higher resolution than ever before. Previous research differentiated JCV into two highly divergent lineages, A and B^6,23,26^. This is occasionally expanded to delineate diversity within the B lineage, known as the B2 “sister clade”^6,26^. To further describe the genetic diversity within the JCV A and B lineages, we created a putative lineage classification system using Autolin (**Fig. 2b**; **Methods**)^27^, which uses greedy maximization to assign lineages to representative phylogenetic nodes. While the limited diversity of the S segment only allows for the calling of major lineages A and B, our new lineage system further designates unique clades within the M and L segments. Specifically, we divide the A lineage into two sub-lineages, A.1 and A.2, with the A.1 lineage further differentiated into 3 minor lineages (A.1.1, A.1.2, and A.1.3). The B lineage is divided into the root B lineage and B.1 sub-lineage, where the previously used B2 “sister clade” primarily falls into the parental B clade. Lineage assignments are robust across the M and L segments, making lineage calling consistent and reproducible.

Upon designating lineages to the JCV sequences, we identified 7 with incongruent lineages across the segments (**Fig. 2c** and **Supplementary Table 2**) and used several complementary statistical analyses (**Methods**) to confirm that mixed calls were the result of reassortment between the segments of different virus lineages. Within our sample set, we found that cross-lineage segment reassortment in JCV was infrequent, representing a ∼1% reassortment frequency with no clear correlation between year, location, or mosquito species, and no detected descendants (**Supplementary Table 2**).

To categorize genetic diversity across the US, we analyzed the spatial distribution of JCV lineages (**Fig. 2d**). Of note, as we were able to sample densely from parts of the Northeast US and the A.2 lineage is largely restricted to the western US (North Dakota, New Mexico), we excluded this lineage from all subsequent phylogenetic analyses. Within our 598 sequences from the Northeast, we found an observable difference between the circulating lineages in New York State and Pennsylvania when compared to Connecticut and Massachusetts. Lineage A sequences from New York State are largely designated as A.1 (29.5%), A.1.2 (35.2%), and A.1.3 (15.6%). The lineage distribution in New York State supports previously observed broad geographic clustering^22^, with increased prevalence of A.1.2 and A.1.3, particularly in western regions of the state. In contrast, lineage A sequences from Connecticut and Massachusetts are almost entirely A.1.1 (94.9%). Due to differences in JCV surveillance and detection methods across states, distribution of B lineages cannot be directly compared (**Methods**). Regardless, heterogeneity in distribution of viral diversity suggests some level of historical spatial restriction of lineage establishment and spread.

### Evolutionary rate of JCV is restricted by annual periods of stasis

Our large JCV genomic dataset (**Fig. 2**) enabled us to estimate the evolutionary rates and patterns of the virus for the first time. To best capture temporal signal across the JCV genome, we removed the reassortant sequences (**Fig. 2c**) and concatenated the S, M, and L genome segments and then tested the temporal signal using a Bayesian phylogenetic approach (**Methods**).

We first analyzed temporal signal and evolutionary rates independently for each of the major A and B lineages (**Fig. 3**; **Supplementary Table 3** and **4).** We estimated that the evolutionary rate for lineage A was 3.57 x 10^-5^ (95% Highest Posterior Density (HPD): 3.09 - 4.11 x 10^-5^) substitutions per site per year (s/s/y) and lineage B was 2.74 x 10^-5^ (95% HPD: 1.85 - 3.72 x 10^-5^) s/s/y. While our JCV evolutionary rate estimates are close to overall the substitution rate for the *Orthobunyavirus* California serogroup (which includes JCV) at ∼6 x 10^-5^ s/s/y^28,29^, they are ∼10-100x slower than the estimates for a related *Orthobunyavirus* (Oropouche virus at ∼1 x 10^-3^ s/s/y^30^), a most distally related *Phlebovirus* (Rift Valley fever virus at ∼3.6 x 10^-4^ s/s/y^31^), and many other mosquito-borne viruses^32–35^, making JCV among the slowest evolving RNA viruses^36,37^ (**Fig. 3**).

**Figure 3.**
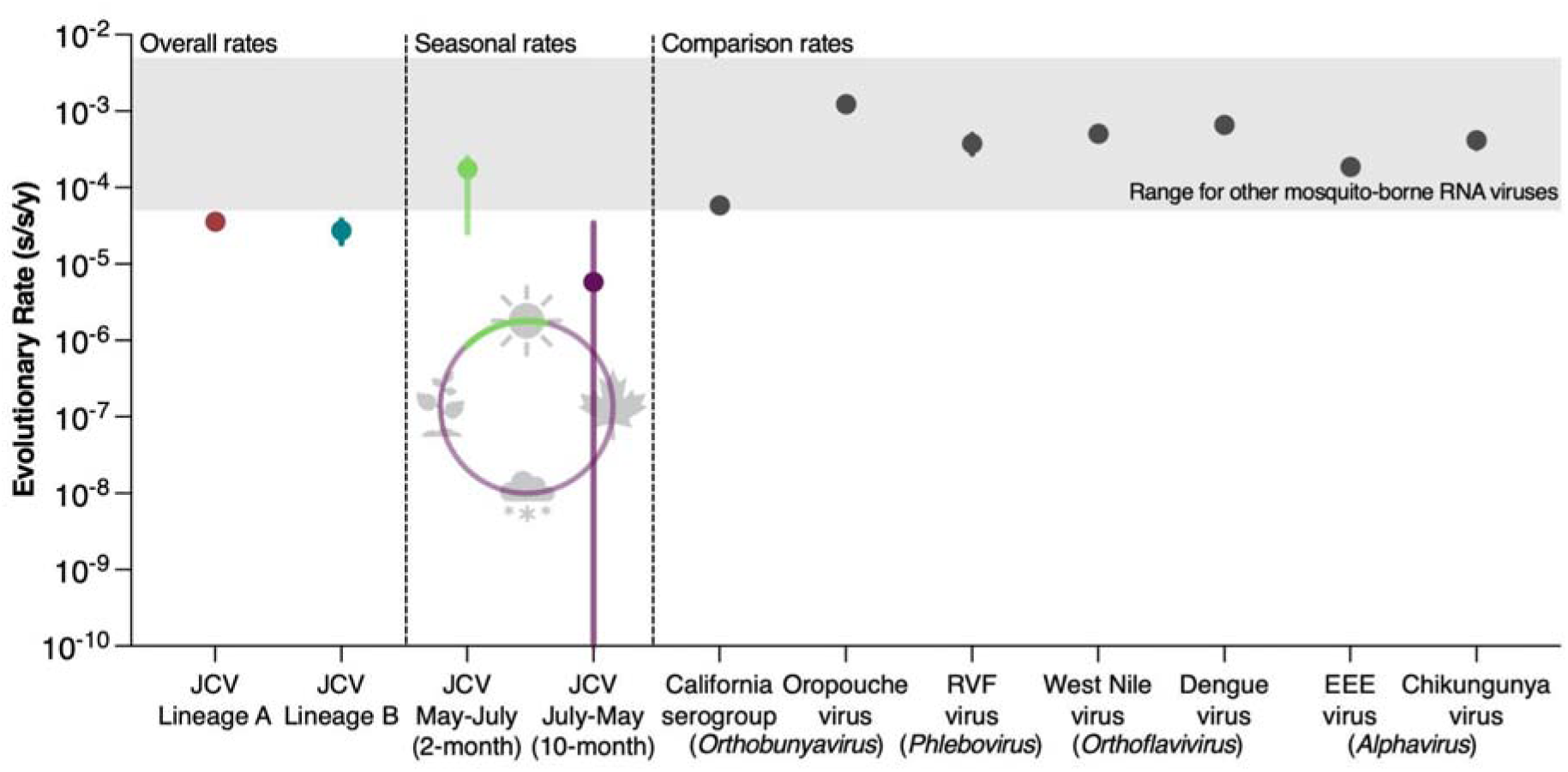
Slow evolutionary rate of JCV relative to other mosquito-borne RNA viruses driven by 10 month cycles of stasis. Point estimates represent median evolutionary rates in substitutions per site per year (s/s/y), with vertical bars showing 95% highest posterior density (HPD) intervals. The ‘overall rates’ are estimated from the concatenated the S, M, and L genome segments with reassortant sequences removed and conducted across the individual phylogenies for JCV lineages A and B. The ‘seasonal rates’ are estimated from an epoch model applied to the substitution rate: (1) each 2-month interval starting at mid-May to mid-July from 1997-2022, (2) each 10-month interval starting at mid-July to mid-May from 1997-2022, and (3) all months prior to 1997 (estimate not shown in the figure). The ‘comparison rates’ were compiled from the literature: overall estimate of closely related *Orthobunyaviruses* (the California serogroup from the Americas^28^ and Oropouche virus (transmitted by midges) during the epidemic in Brazil^30^), a distally related *Phlebovirus* (Rift Valley fever (RVF) virus from Africa^31^), and other mosquito-borne viruses of the *Orthoflavirus* (dengue virus [serotype 1] from a global dataset^35^ and WNV lineage 1a from North America^33^) and Alphavirus (EEEV from the eastern US^32^ and the chikungunya virus Asian lineage^34^) genus. The grey shaded area between 5 x 10^-5^ and 5 x 10^-3^ s/s/y is the previously known expected evolutionary rate range for mosquito-borne RNA viruses^36,37^.

We hypothesized that the evolutionary rate of JCV may be restricted by how the virus over-winters. Single-brood (univoltine) mammal-biting *Aedes* mosquitoes are thought to be important for trans-seasonal JCV persistence^6,38^ in part from field data suggesting that the virus can over-winter in eggs via transovarial transmission^39–43^. To test our hypothesis that JCV evolves more slowly while over-wintering, we employed two different epoch models to the substitution rate, jointly for lineages A and B (**Methods**). First, we defined the epochs as (1) a 2-month interval from mid-May to mid-July corresponding with univoltine *Aedes* mosquitoes emergence^6^ and (2) a 10-month interval for the rest of the year from 1997-2022 (**Fig. 3**). We estimated that the median JCV evolutionary rate for the 2-month interval was 1.72 x 10^-4^ s/s/y (95% HPD: 2.59 x 10^-5^ - 2.23 x 10^-4^ s/s/y), which is similar to the range of other mosquito-borne RNA viruses, while the 10-month interval was 5.81 x 10^-6^ s/s/y (95% HPD: 8.83 x 10^-15^ - 3.44 x 10^-5^ s/s/y) with a posterior mass close to 0 (**Extended Data Fig. 2**). When we extend the epoch range to a 4.5-month interval from mid-May to the end of October corresponding to when multivoltine mosquitoes are generally active in Connecticut, the median evolutionary rate decreased to 6.53 x 10^-5^ s/s/y (95% HPD: 1.38 - 8.2 x 10^-5^ s/s/y) with the remaining 7.5-month interval rate remaining low at 6.57 x 10^-6^ s/s/y (95% HPD: 7.38 x 10^-14^ - 4.95 x 10^-5^ s/s/y), indicating that JCV evolves the fastest during the 2 months from mid-May to mid-July. Overall, we show that JCV evolves at a similar rate to other mosquito-borne RNA viruses (∼10^-4^ s/s/y) for ∼2 months of the year from mid-May to mid-July, but then its evolutionary rate approaches zero for the other 10 months, likely while overwintering in dormant mosquito eggs and suggesting the there is little to no viral replication during the period.

### Phylogeographic reconstruction of JCV in the Northeastern US

Having identified genetically distinct JCV lineages in the Northeastern US (**Fig. 2**), we sought to estimate when the major introductions occurred into the region. As with our evolutionary analyses above, we removed the reassortant sequences and concatenated the S, M, and L genome segments of samples collected from the Northeast prior to conducting discrete phylogeographic analyses (**Methods**). We assigned each sequence to the state in which it was collected and reconstructed the overall introduction times and likely transitions between states independently for both A and B lineages (**Fig. 4**). Because Connecticut’s mosquito surveillance program has amassed the most comprehensive collection of JCV-positive mosquito pools in the US (597 total, 1997-2022)^6^, of which we sequenced 84% of all positive samples, we centered our analyses on when, and how often, JCV was introduced into the state.

**Figure 4.**
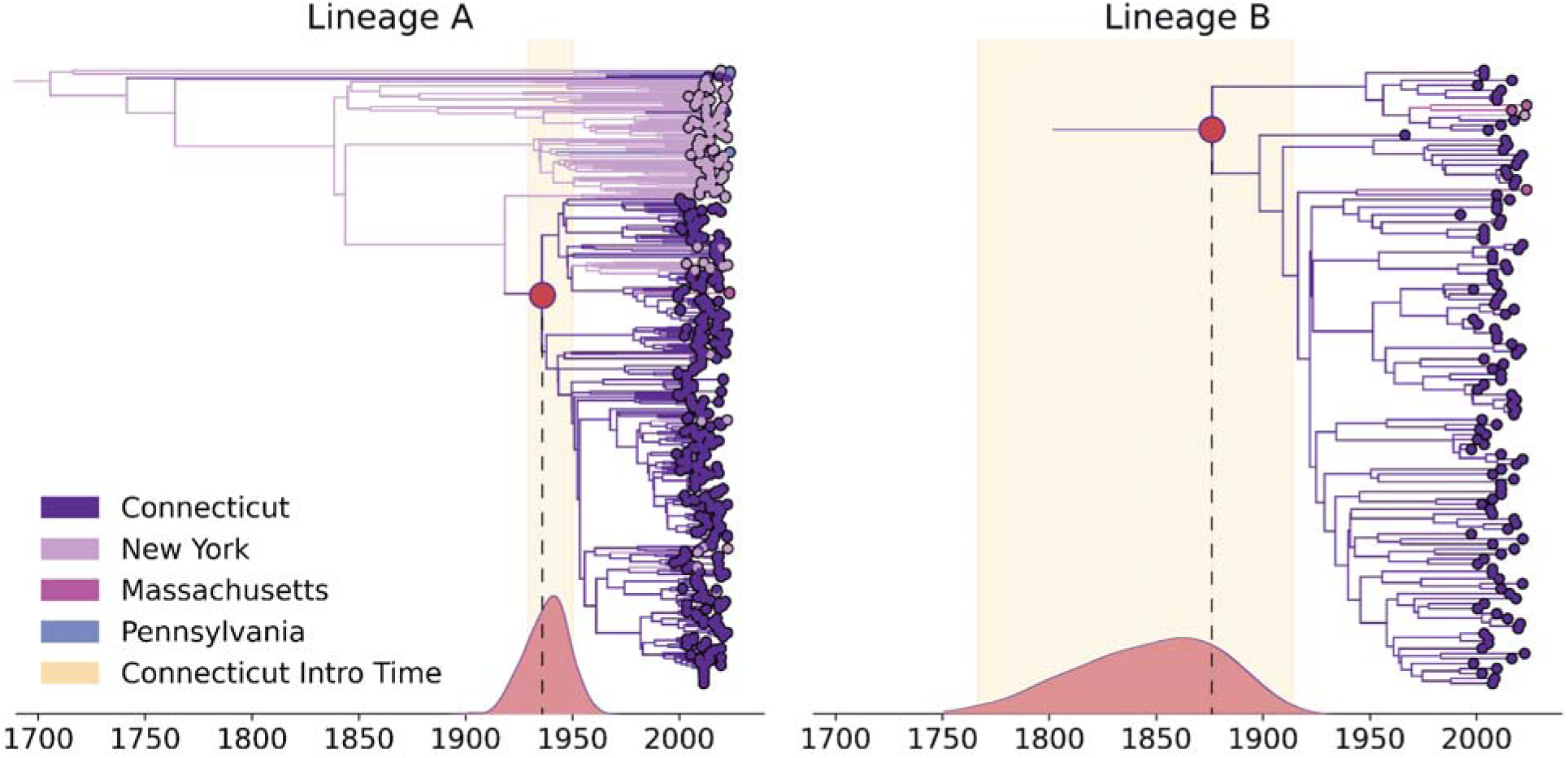
Discrete phylogeographic reconstruction of JCV lineages shows long-term persistence in Connecticut. Time-scaled phylogenies of concatenated S, M, and L genome segments for lineage A (left) and lineage B (right), as inferred using a Bayesian phylogeographic analysis that employs an asymmetric forward-in-time CTMC model. Tip points are colored by sampling location, with internal branches and nodes colored according to the most probable inferred state on the maximum clade credibility (MCC) tree. Red dots denote the most basal Connecticut node. Yellow shading area indicates the corresponding 95% highest posterior density (HPD) interval for the timing of this node. The kernel density estimation of the node’s timing based on 1000 posterior samples is plotted along the x-axis in red.

Our time-resolved, discrete phylogeographic analysis of JCV lineage A estimates the time of the most common recent ancestor (tMRCA) of all of the Northeastern sequences to be in the early 1700s (95% HPD: 1659-1788; **Fig. 4**). Based on our sequences, the surviving descendents of the lineage A ancestor persisted in New York for ∼200 years before they were introduced into Connecticut. The limited availability of lineage B sequences outside of Connecticut makes it difficult to estimate the tMRCA for the region, although it has likely been circulating for over one hundred years.

We detected at least 4 separate introductions into Connecticut (3 lineage A, 1 lineage B; **Fig. 4**) The oldest tMRCA of a lineage A clade in Connecticut was in the early 1900s (95% HPD: 1920 to 1947) and represents the first inferred JCV introduction in Connecticut from New York. The subsequent expansion of lineage A.1.1 in Connecticut led to at least 10 re-introductions into New York and onward spread into Massachusetts. The tMRCA of lineage B in Connecticut was in the late-1800s, though with a very broad range of inferred dates (95% HPD: 1759 to 1903). This uncertainty likely reflects the lack of lineage B sequences from outside of Connecticut to provide context, making it difficult to estimate introduction times into the state. The separate B and B.1 lineage clades in Connecticut (**Fig. 2b,d**) could be the result of multiple introductions that occurred in the 1900s, closer to our estimated lineage A introduction date. Despite this uncertainty, it is clear that JCV has a long history of circulation in the Northeast, dating back at least 100 years in Connecticut **(Fig. 4)**.

### Phylodynamics of mosquito traits infers mechanism of JCV persistence

While we show that JCV has persisted in Connecticut for at least a century (**Fig. 4**), the mechanisms for long-term local maintenance of the virus are not completely understood. JCV infects a diversity of mosquitoes and is not reliant on any one species^6,7,42–44^. From the active JCV surveillance system in Connecticut, we detected the virus in 597 mosquito pools representing 26 different mosquito species from 1997 to 2022 (**Fig. 5a**). Surveillance data from Connecticut support the role of univoltine *Aedes* mosquitoes for JCV maintenance and persistence^6,7,44,45^. We show that univoltine species emerge from vernal pools from mid May to early June, rapidly expanding in population size to peak in the early summer months, corresponding to when JCV is often detected (**Fig. 5b**, left). In contrast, multi-brood (multivoltine) mosquitoes demonstrate slower emergence, with a more sustained population throughout the summer persisting into the fall. JCV infection rates in univoltine mosquitoes are typically higher than multivoltine species (**Fig. 5b**, middle). Combined, our estimated vector index, a measure of mosquito-borne pathogen risk to humans^46^, shows a higher risk in univoltine mosquitoes, especially during the early season (**Fig. 5b**, right). These differences between mosquito phenologies clarify key aspects of the complex JCV transmission cycle, highlighting unique roles of univoltine and multivoltine species in the seasonality of the virus. We hypothesize that early season transmission is driven by the emergence of infected univoltine mosquitoes, with transmission maintained throughout the summer by the extended emergence of multivoltine species.

**Figure 5.**
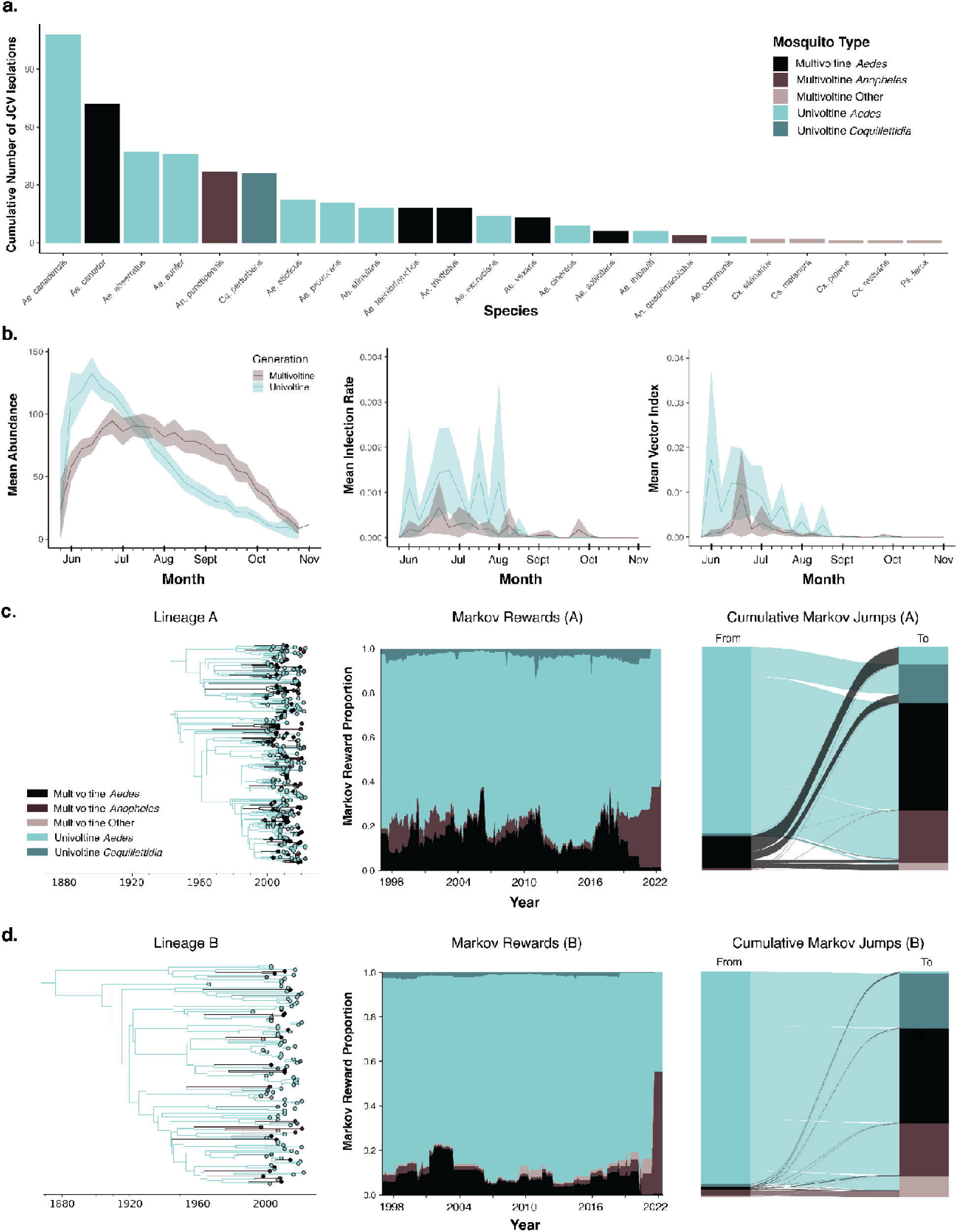
Ecological and phylogenetic analyses reveal that univoltine *Aedes* mosquitoes are responsible for JCV persistence. (**a**) Cumulative detections of JCV by mosquito species across the full surveillance period, 1997-2022. Bars are colored according to the genus and generation type of mosquito of the listed species; (**b**) Mean abundance, pooled infection rate, and vector index of univoltine and multivoltine mosquitoes in Connecticut, averaged per trap per generation group per epi-week from 1997-2022; (**c**) Discrete trait phylodynamic analysis of lineage A JCV in Connecticut mosquitoes. Left: Bayesian maximum clade credibility (MCC) tree with tip points colored by mosquito pool of origin, with internal branches and nodes colored by most probable mosquito type. Middle: Markov rewards proportion showing the expected fraction of lineage A time spent in each mosquito group along the phylogeny (1997-2022). Right: Alluvial plot of cumulative Markov jumps, with bars on the representing the category jumped from and bars on the right representing the category jumped into; (**d**) Discrete trait phylodynamic analysis of lineage B JCV in Connecticut mosquitoes, displayed as in panel (**c**). Left: MCC tree with tip and node coloring by mosquito type. Middle: Markov rewards proportion. Right: Alluvial plot of cumulative Markov jumps.

By integrating mosquito identity (generation type and genus) associated with each JCV sequence from Connecticut as a discrete trait within our phylodynamic framework, we reconstructed how the virus persists in different mosquitoes (**Fig. 5c-d**). The comprehensive surveillance data in Connecticut, available statewide at multiple time points per season (**Methods**) facilitated our highly complete genomic data, containing nearly all (83.9%) JCV-positive mosquito pools actively collected since 1997. As a result, phylogenetic data, and corresponding ecological metadata, are representative of JCV circulation in Connecticut mosquitoes and do not require further subsampling.

Our phylodynamic analyses show that the majority of the ancestors of our JCV sequences (for both A and B lineages) were inferred to be univoltine *Aedes* mosquitoes (**Fig. 5c-d**, left). Specifically, we quantified across the lineage A and B phylogenies that JCV spent 78.9% and 90.3% of time, respectively, in univoltine *Aedes* mosquitoes (Markov rewards; **Fig 5c-d**, middle). Comparatively, only 57.1% of JCV-positive mosquito pools were from univoltine *Aedes* (**Fig 5a**), indicating that the role of this mosquito group in JCV persistence is even greater than what can be inferred from surveillance alone. We also quantified that 83.6% to 94.4% (lineages A and B, respectively) of transitions between mosquito groups originate in univoltine *Aedes*. (Markov jumps; **Fig. 5c-d**, right). We detected some persistence and spread from multivoltine *Aedes*, and to some extent, multivoltine *Anopheles* (**Fig. 5c-d**), but at a lower rate than their detections in mosquitoes (**Fig 5a-b**). Our analysis demonstrates that univoltine *Aedes* mosquitoes are necessary for long-term virus persistence, providing further evidence for overwintering in these mosquitoes, with multivoltine species potentially acting as secondary vectors in the JCV transmission cycle.

### Ecological determinants of JCV dispersal

While many mosquito vectors may have roles in short-term JCV transmission, a few key univoltine *Aedes* species are likely responsible for long-term persistence (**Fig. 5**). These boreal, or snowpool, *Aedes* mosquitoes emerge in large broods in the late spring and early summer, seeking white-tailed deer and other mammals for bloodmeals^44,47^. Over the course of their short lifespans, these mosquitoes produce multiple clutches of eggs that remain dormant through overwintering, hatching only after surviving a freeze-thaw cycle in the following season^44,47^. If these species are responsible for maintaining JCV, they provide limited opportunities for virus dispersal. These ecological constraints likely limit lineage-level dispersal, shaping the spatial structure of JCV transmission. As previously described^6,7,44,45^, we observed differences in the spatial distributions of JCV lineages A and B (**Fig. 6a-b**), with a higher proportion of lineage A detections in southwestern Connecticut and lineage B more evenly distributed across the state. Our reconstructions of the effective population size for both lineages indicate that viral populations are similar in size and have largely remained stable across the surveillance periods (**Extend Figure 3a**).

**Figure 6.**
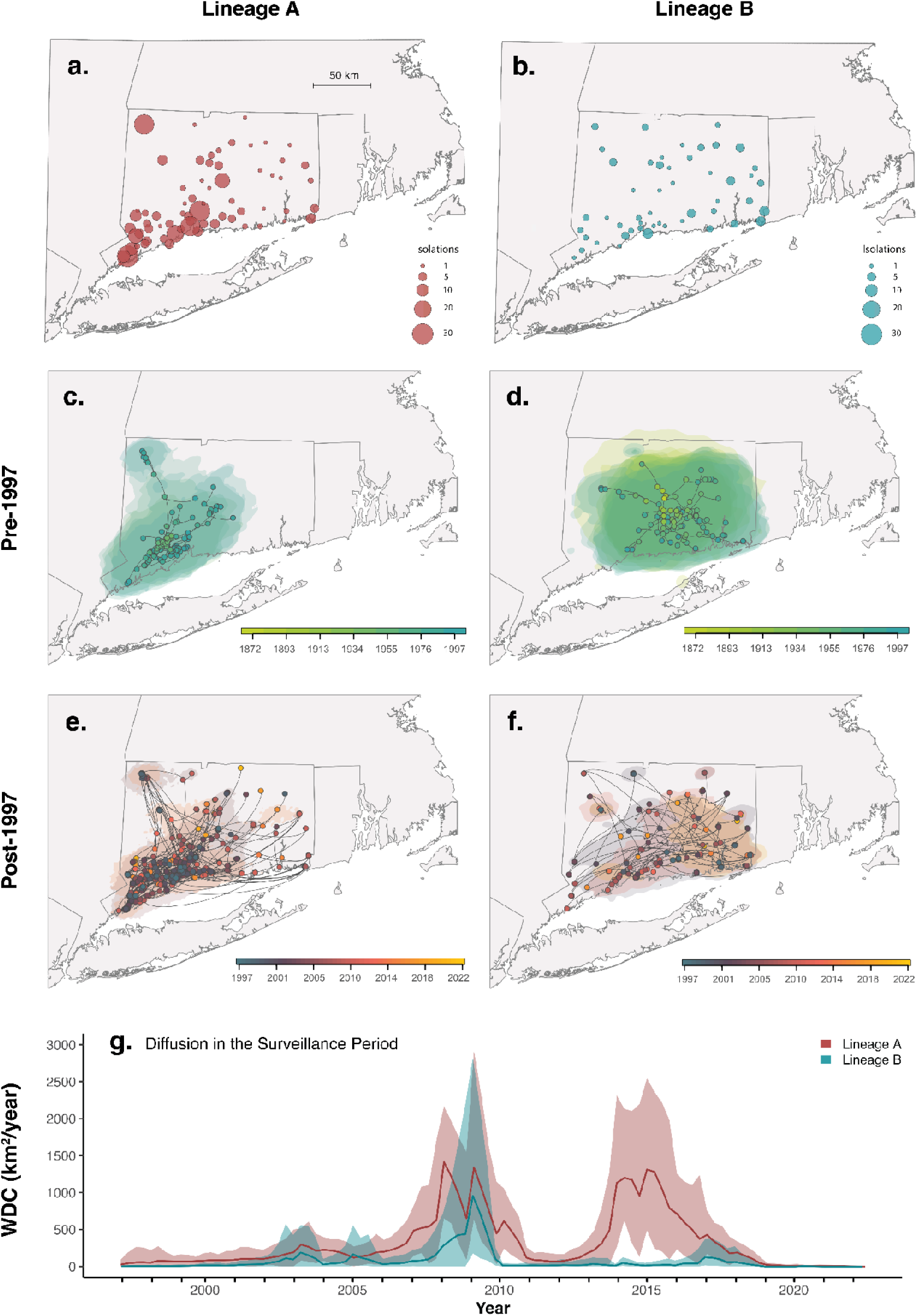
Phylogeographic analysis shows patterns of slow long-term JCV dispersal intermixed with short bursts of diffusion in Connecticut. (**a - b**) Isolation of lineage A (left) and B (right) JCV across traps in Connecticut. Each point represents a distinct trapping location, with the size of the point scaled to reflect the total number of virus isolations at each site. (**c - d**) Continuous phylogeographic reconstruction of viral movements for major lineages A (left) and B (right) prior to the start of surveillance in 1997; (**e - f**) Continuous phylogeographic reconstruction of viral movements for major lineages A (left) and B (right) in the period of seasonal surveillance, post-1997. Polygons represent inferred 80% highest posterior density (HPD) regions colored according to inferred occurrence time. Curved transmission lines indicate the direction of viral movement, with counterclockwise lines indicating a movement from origin to destination; (**g**) Weighted diffusion coefficient (WDC, km^2^/year) of JCV lineages over time. Lines indicate the mean value of the WDC, while shaded regions indicate the 95% HPD region of this estimate.

To further investigate how the JCV transmission cycle impacts lineage dispersal, we performed independent continuous phylogeographic reconstructions of major lineages A and B in Connecticut (**Fig. 6**). As opposed to discrete phylogeographic reconstruction (**Fig. 4**), continuous phylogeography allows us to link specific coordinates to our sequences, which can evolve along the branches of the tree to make precise spatiotemporal inferences for calculating dispersal capacities and comparing diffusion patterns among lineages (**Methods**)^19,48^. To account for potential uncertainty in the period before the Connecticut mosquito surveillance program began in 1997, we divided our phylogeographic analyses into two distinct time periods. All movements pre-1997 represent historical dispersal, a period from Connecticut when we have limited sequencing (2 sequences, 1966 and 1992), and all patterns are based on inferred ancestral nodes of contemporary viruses (therefore without including any potential extinct branches; **Fig. 6c-d**). Post-1997 describes the period of surveillance and dense sequencing, or contemporary dispersal, when we can reconstruct the movements with higher confidence (**Fig. 6e-f**). Our reconstructions of the historical dispersal patterns reveals that lineage A was likely introduced into southwestern Connecticut (**Fig. 6c**) while lineage B was likely introduced into the central part of the state (**Fig. 6d**) during the late 1800s or early 1900s. For the ancestors of both lineages, we estimate that historical dispersal was restricted to ∼30-50km radius around the inferred introduction locations. After 1997, the contemporary dispersals of both lineages reached all regions of the state; however, the dispersal of lineage A remained concentrated in the southwestern region (**Fig. 6e**) while lineage B appeared to be less geographically restricted (**Fig. 6f**). Thus, the distinct spatial distributions of lineages A and B detected from the surveillance system (**Fig. 6a-b**) may be the result of different introduction locations and slow dispersal (founder effect). With time, the lineage A and B distributions may become completely intermixed or remain separated due to differences in ecological preferences or inter-lineage competition.

To determine if environmental factors were associated with lineage dispersal dynamics, we performed landscape phylogeographic analyses for each lineage, testing the potential impacts of features such as elevation, precipitation, and temperature on lineage dispersal (**Methods**). We found that inferred JCV lineages tended to avoid circulating in croplands and areas of increased elevation (**Supplementary Table 6**), as these are not frequent habitats for mosquitoes like *Ae. canadensis* that develop in vernal, leaf-lined woodland pools^6,47^. We did not, however, find statistical support for associations between the environmental factors tested here and the diffusion velocities of inferred JCV lineages (**Supplementary Tables 7**). We cannot rule out the possibility that other, non-tested, environmental factors and/or a combination of various factors explain the differences in lineage spread.

Overall, our analyses suggest that JCV is typically a very slow-moving virus with intermittent bursts of spread. We evaluated continuous phylogeographic spread by the weighted diffusion coefficient (WDC), a two-dimensional measure of virus “diffusivity”^49,50^. We estimated the historical WDC of lineage A to be 32.20 km^2^/year (95% HPD: 27.73-37.47) and lineage B at 19.33 km^2^/year (95% HPD: 13.25-26.99; **Supplementary Table 5**), while they respectively increase to 57.22 km^2^/year (95% HPD: 50.03-64.17) and 62.38 km^2^/year (95% HPD: 49.12-78.31) during the post-1997 contemporary dispersal period (**Supplementary Table 5**). These rates are on the same scale as our previous estimates for the tick-borne Powassan virus (∼130 km^2^/year)^50,51^, but far slower than other mosquito-borne viruses such as WNV and yellow fever virus^50,52,53^. The yearly WDC estimates, however, were variable, peaking at a maximum of ∼1,000 km^2^/year in 2008-2009 for both lineages and again for lineage A around 2015 (**Fig. 6g**; **Extended Data Fig. 3**). These long-range lineage dispersal events would cover approximately 7% of the total 13,023 km^2^ area of Connecticut in one year^54^. Although long-term JCV lineage spread is slow compared to other mosquito-borne viruses, these periodic bursts of diffusion may introduce the virus into new areas of the state.

To further test the impact of ecological covariates on the temporal dynamics of JCV diffusion, we fit a Bayesian generalized additive model (GAM) to posterior WDC data frames for each lineage over time (**Methods**; **Extended Data Fig. 4**). Estimates of WDC are emergent, model-dependent quantities that combine effects of phylogenetic structure, sampling, and spatial heterogeneities^55^. While this model cannot directly demonstrate mechanisms of lineage diffusion, it can be used in a descriptive capacity in support of our mosquito surveillance and phylodynamic analysis to understand the ecology of JCV. To account for variation in tree topology and uncertainty in virus dispersal, our model included the impact of the number of phylogenetic branches per window as a smoothing term, capturing possible biases inherent to WDC calculation and allowing for uncertainty across posterior trees. For lineage A, annual diffusion rates were negatively associated with precipitation (−0.73, 95% HPD: [−0.75; −0.70]; **Extended Data Fig. 3**). The impact of branch number on WDC was strong (3.42, 95% HPD: [3.04; 3.83]), indicating variation in diffusion that reflects episodic bursts as opposed to continuous spread. In contrast, lineage B demonstrated distinct sensitivities to several of the selected variables. Precipitation (2.43, 95% HPD: [2.32; 2.53]) and multivoltine mosquito abundance (0.48, 95% HPD: [0.42; 0.53]) were positively associated with WDC, while univoltine mosquito abundance was negatively associated (−0.20, 95% HPD: [−0.23; −0.17]). When this category is further expanded into generation-genus groups, lineage B displays the strongest positive association with multivoltine *Anopheles* mosquitoes (1.10, 95% HPD: [1.05; 1.14]), while multivoltine *Aedes* have a significantly lower contribution (0.38, 95% HPD: [0.34; 0.42]). For lineage B, the smoothing term for phylogenetic branching was weaker (0.72, 95% HPD: [0.23; 1.19]), indicating more stable diffusion over time.

Together, these results support the observed localized persistence of JCV. Coupled with our previous observations regarding the evolutionary rate and maintenance in univoltine *Aedes* mosquitoes, we hypothesize that historical and local ecological processes, rather than ongoing rapid movement, produced observed lineage heterogeneity. Our findings further suggest that transient spillover into multivoltine *Anopheles* populations (**Fig. 5**) during wet and warm periods may facilitate short bursts of geographic expansion, potentially elevating seasonal human infection risk.

## Discussion

JCV is a prevalent yet underrecognized arbovirus of public health concern, necessitating further study into its evolution and ecology. To address this, we developed an amplicon-based sequencing approach and generated 658 complete JCV genomes, expanding available genomic data nearly 17-fold. With these new data, we provide the first estimate of the JCV evolutionary rate at ∼3 x 10^-5^ s/s/y, defined by prolonged periods of stasis, making it among the slowest evolving RNA viruses^36,37^. When the univoltine *Aedes* mosquitoes are active during the early summer months, JCV evolves at rates typically estimated for other mosquito-borne viruses (10^-4^ to 10^-3^ s/s/y); however, its evolution nearly stops for the rest of the year, suggesting that the virus does not replicate while overwintering in mosquito eggs. Despite this, JCV has accumulated substantial genetic diversity through centuries of evolution in the Northeastern US. We further classified the previously defined A and B lineages^6,23,26^ into several distinct sublineages to aid future tracking of the virus.

We estimated the precise patterns of JCV spread and transmission by sequencing 84% (500/597) of the JCV-positive mosquito pools collected from 1997-2022 through an active surveillance program in Connecticut^6^. Our phylogeographic inference suggests that JCV was introduced several times in Connecticut, with the primary introductions of lineages A and B occurring in the mid 1800s to 1900s. Despite frequent isolation from a broad diversity of mosquito species, our phylodynamic discrete trait analyses identified univoltine *Aedes* mosquitoes as the primary driver of long-term local persistence in Connecticut. We hypothesize that the dependence on these single-brood mosquitoes limits its local diffusion rate to ∼30-60 km^2^/year, which is more similar to slow-moving tick-borne viruses than other mosquito-borne viruses^50^. In contrast, we show that JCV transmission by multivoltine mosquitoes was associated with short-term bursts in viral diffusion. Our study demonstrates the extent to which dense sequencing and phylodynamics can rapidly advance our knowledge of the evolutionary history and complex life cycle of a previously understudied virus.

Building on this foundation, we propose that JCV persists and spreads in Connecticut through two ecological cycles that sustain its trans-seasonal maintenance and within-season diffusion (**Fig. 1**). The “maintenance cycle” is primarily driven by univoltine *Aedes* species, such as *Ae. canadensis*, *Ae. aurifer*, *Ae. abserratus, Ae. strictus,* and *Ae. provocans*. These mosquitoes emerge in the late spring and early summer, with some individuals already infected with JCV via transovarial transmission^39–43^. Although transovarial transmission is usually not particularly efficient, JCV transmission to mammals subsequently aids the early season amplification of infected mosquitoes. White-tailed deer are a bloodfeeding source for many of these univoltine *Aedes* species in the region^6,20,56^ and are recognized as the principal JCV amplification host based on experimental infections and high seroprevalence rates^12,57–61^ (though other deer species may have a similar role in other regions^6^). Infected females then oviposit JCV-infected eggs that enter diapause, overwintering for ten months before reinitiating transmission the following year^44,47^. These episodic evolutionary dynamics punctuated by annual periods of apparent stasis explain both the slow rates of JCV evolution and spread.

We further hypothesize that the enzootic maintenance of JCV described above is complemented by the “diffusion cycle” (**Fig. 1**) facilitated by multivoltine *Anopheles* mosquitoes, including *An. punctipennis* and *An. quadrimaculatus*, that remain active throughout the summer through successive broods of offspring^44,47^. As these species also feed on white-tailed deer^56^, JCV-infected deer are likely the connection points between the two cycles. The extended seasonal activity of multivoltine *Anopheles* mosquitoes thus increases opportunities for onward enzootic transmission, enabling JCV to at least temporarily spread beyond univoltine *Aedes* habitats. While most of these long-distance movements ultimately result in dead-ends for transmission at the end of the season (possibly due to inefficient virus transovarial transmission prior to overwintering as un-bloodfed adults), such episodic bursts may correspond with increased risk of human infection^38^. Future study is critical to further understand the role of our proposed diffusion cycle and to evaluate whether there are entomological indices that can be used to forecast JCV transmission. Targeted seasonal sampling of mosquito vectors and hosts could clarify the timing and extent of diffusion events, improving prediction of human JCV risk.

Our phylogeographic analyses estimate that JCV has a long-established history in the Northeastern US. We show that JCV Lineage A was likely introduced in the region during the early 1700s indicating that the virus has been locally maintained for nearly three centuries. In at least some parts of the Northeast, likely including New York, this suggests JCV persisted through the dramatic decline of white-tailed deer population in the late 1800s driven by habitat loss and overhunting^57,62^. In Connecticut, JCV appears to have become established later, around the early 1900s, coinciding with forest regrowth and the subsequent rebound of deer populations^7,56,62^. Given its long history in the Northeast and widespread establishment, JCV likely has very old roots in North America and persisted through several major ecological changes. Indeed, JCV is very diverse, and the species now officially includes the related South River virus from New Jersey and Jerry Slough virus from California^63^. One estimate places the common ancestor of JCV and its relatives in North America at more than 1,000 years ago^29^.

While we significantly increased the amount of JCV genomes, we still lack sequencing data from diverse locations across the continent that could provide more precise timings on its emergence and spread over ecological gradients and throughout long periods of time. Specifically, there is limited JCV genomic data from key Midwestern states such as Minnesota, Michigan, and Wisconsin that report the highest rates of human cases^17,20^. These biases restrict our ability to assess national transmission dynamics and make broader inferences regarding viral diversity and diffusion patterns. The limited availability of lineage B sequences outside of Connecticut, potentially driven by differences in molecular detection methods, constrained our reconstruction of its evolutionary history. Although mosquito surveillance efforts have increased in the Midwest^43^, more targeted sequencing is critical to capture the full spectrum of regional ecological variation. For example, in North Dakota, JCV has been primarily associated with *Aedes vexans*, a multivoltine species^64^, suggesting that the roles of mosquito vectors may differ across landscapes. Further, research is still needed to disentangle which univoltine *Aedes* species are the most important JCV maintenance, even in the Northeast. Mosquito surveillance in Connecticut (and New York) typically starts around June 1, which likely misses the earlier emergence of vernal pool mosquitoes - especially *Ae. provocans*. When *Ae. provocans* are collected in Connecticut from early mosquito pools, there is a high odds ratio that JCV will be isolated^44^. As is, our JCV data are probably skewed toward *Ae. canadensis* that typically emerge in June and are more likely to be captured by surveillance. The type of spring can also influence the dynamics of univoltine mosquito populations, with heavy rainfall favoring species like *Ae. strictus* that are associated with floodplains.

Our study also cannot account for the role of white-tailed deer in JCV maintenance and spread, as spatial data on deer populations are limited and JCV investigations of deer are sparse^6,7,21^. While deer are likely key amplifying hosts, and recent work suggests univoltine mosquitoes initiate seasonal transmission of JCV in deer^62^, we could not directly assess how host movements interact with mosquito dynamics to shape broader transmission patterns. Addressing these limitations will require expanded field studies and genomic sampling, particularly in regards to white-tailed deer. Coordinated JCV testing in deer and cross-state data sharing using wildlife surveillance standards would improve our understanding of virus circulation^65^.

Our study provides a critical foundation for understanding the viral diversity, transmission dynamics, and ecological persistence of JCV. We demonstrate how combining phylodynamic and ecological frameworks can uncover patterns of arbovirus transmission and spread otherwise obscured by complex ecologies, sparse surveillance, and limited research support. Our results not only clarify the ecological roles of univoltine and multivoltine mosquitoes in sustaining and amplifying JCV transmission, but also offer a scalable model for investigating other understudied arboviruses. Continued investment in sequencing and targeted vector surveillance will be essential to capture the full extent of diversity of emerging viruses. These data, when paired with mixed phylodynamic analyses, enable us to infer long-term circulation patterns, dispersal processes, and ecological interactions that drive virus maintenance and spread. As climate change, habitat fragmentation, and shifting host distributions alter the ecology of arboviruses^19^, analytical approaches like ours will be increasingly important. This work reinforces the power of integrating evolutionary and ecological data to disentangle complex virus transmission cycles, underscoring the need to apply integrated phylodynamic approaches to emerging and understudied viruses.

## Methods

### Sample Collection and Assessment

All samples from Connecticut were collected as part of the Connecticut Agricultural Experiment Station’s (CAES) mosquito surveillance program, which has detected JCV since the start of its operation in 1997^45^. While originally targeting 37 trapping locations, the program has repeatedly expanded, reaching 73 sites in 2000, 91 from 2001-2015, 92 from 2016-2019, and 108 from 2020-2022. A full list of JCV isolations and their corresponding trapping locations can be found at: https://github.com/grubaughlab/JCV-genomics. The program is ongoing, with available surveillance data from our sequencing period, 1997-2022. Each location accommodates one CDC CO_2_ baited light trap and one gravid trap, which are set in the late morning and collected the following day. The CAES conducts surveillance from June to October each year and sets traps at each site approximately every 10 days, which increases to twice a week if recently trapped mosquitoes test positive for WNV or EEEV. Collected mosquitoes are pooled by species, location, and date and tested for arbovirus infection by cell culture and RT-PCR methods as previously described^7^. This program not only provided all JCV samples from Connecticut, but also its associated surveillance data, as described in the section below (Surveillance and Seasonal Modeling).

Samples provided by CAES also included 60 JCV-positive mosquitoes from North Dakota, collected from 2003-2006^64^. Traps with CO_2_ light traps were established in Cass and Williams counties, with variable sampling at a variety of trapping locations. Pooling and testing protocols for arboviruses, utilizing cell culture and RT-PCR, were in line with previously described CAES methods^64^.

JCV-positive mosquito pools from New York were also collected as part of a statewide surveillance program, with collection periods ranging from 2003-2022^22^. In a similar system to Connecticut and North Dakota, CDC CO_2_ light traps were deployed and individuals from the same species were pooled into groups of 10-60 mosquitoes prior to testing^66^. All RT-qPCR screening for JCV in New York State (NYS) was performed using a primer and probe set specific to lineage A^22^, which may lead to under-detection of other lineages. However, NYS also tests pools using a generic orthobunyavirus primer set to compensate for possible deficits^22^. Amplified JCV products from RT-qPCR were amplified using a single forward and reverse primer pair in the UTR regions of each segment^22^ and sequenced by the Wadsworth Center Advanced Genomic Technologies Core to confirm sample identity.

Mosquito pools from Massachusetts were provided by the University of Massachusetts Amherst^67^. Mosquito surveillance throughout the state is conducted by State Reclamation and Mosquito Control Boards (SRMCB) across 11 regions in the Commonwealth of Massachusetts^68^. A variety of different trapping methods are employed, including larval dip, CO_2_, light, and gravid traps. The testing of mosquitoes, pooled by trap, species, and date, are performed at the regional level.

In Pennsylvania, mosquito monitoring and control is primarily performed by the Mosquito Borne Disease Control Program, an organization under the state’s Department of Environmental Protection (DEP)^69^. Mosquito surveillance is conducted from April to October through the use of gravid and dry ice-baited traps. The DEP is responsible for mosquito species identification, pooling, and JCV testing by RT-PCR^70^.

While extracted samples were provided from New York, Massachusetts, Michigan, and Pennsylvania, we performed our own nucleic acid extraction on all samples from Connecticut using the MagMAX viral/pathogen nucleic acid isolation kit (Thermo Fisher Scientific) and the Thermo Fisher protocol with reduced extraction volumes (100 uL).

To assess samples for virus concentration, we employed a probe-based qPCR assay using the Luna Universal Probe One-Step RT-qPCR kit (New England Biolabs). Primers and probes were previously designed by CAES, targeting JCV’s small segment. These are CAES_JCV154_F (TAATGCAGCAAAAGCCAAAG), CAES_JCV307_R (AAGCCGATGGATGGTAAGAT), and CAES_JCV181_P (CGCTCGTAAACCGGAGCGGA). All primers and probes were diluted to 10 uM working stocks. The qPCR protocol was validated using a positive control gene fragment (gBlock) of the targeted region on the small segment, tested in a triplicate dilution series. A positive control of 1 x 10^3^ copies per uL was included in every qPCR run to ensure consistency across plates, in addition to a negative template control to check for potential contamination. The full qPCR protocol, with full primer, probe, and gene block sequences, along with thermocycler conditions, is available at protocols.io^71^.

### Sequencing Primer Design

In order to capture all three segments of the JCV genome, we employed a multiplexed approach to primer design by combining previously developed JCV segment-specific primers and an internal set generated by PrimalScheme. To maximize genome coverage, we developed our own highly multiplexed PCR primers for the medium and large segments using PrimalScheme^25^. Based on publicly available sequences, we chose to make two separate primer sets for the highly divergent A and B lineages, each forming 800 bp amplicons. Because the A and B primers share identical binding sites, they can be mixed together by pool during the sequencing protocol to cover JCV diversity across lineages. The final primer set consists of segment specific small segment UTR primers, segment specific medium and large UTR primers, and internal A and B lineage primers for the medium and large segments. A full list of JCVSeq sequencing primers can be found in **Supplementary Table 1.**

### Targeted Amplicon-Based Sequencing (JCVSeq)

In order to process the high volume of historical JCV samples, we developed an amplicon-based sequencing method, JCVSeq. We used an adapted Illumina COVIDSeq protocol for all amplicon sequencing, substituting the primers in the RUO COVIDSeq kit for our novel JCV A and B set. Individual A and B primers were diluted in nuclease-free water to a concentration of 20 uM and combined at equal volumes into two separate pools, AB1 and AB2, to prevent overlapping primers in opposing pools from producing truncated products. Due to JCV’s segmented genome, and lack of internal primers on the small segment, the cDNA synthesis and PCR amplification steps of the COVIDSeq protocol were modified and combined. This “hybrid” one-step protocol allows for the simultaneous amplification of the small segment and generation of medium and large segment amplicons. The resulting one-step protocol utilizes IPM, FSM, and RVT to generate cDNA under consolidated thermocycler conditions before proceeding with amplicon generation. Negative template controls were included for each run during the one-step cDNA synthesis/amplification stage. The full sequencing protocol, with full primer sequences, amplicon pools, and thermocycler conditions, is available at our JCVSeq protocols.io. Final pooled libraries were sequenced at the Yale Center for Genome Analysis, with paired-end 150bp reads generated at 1 million reads per sample on an Illumina NovaSeq 6000.

Consensus genomes were generated using a modified version of an iVar-based^72^ analysis pipeline that was originally developed for dengue virus^73^, which maps raw Illumina reads against multiple representative virus references. The pipeline utilized separate reference sequences for each segment of both JCV A (GenBank accession numbers HM007356.1, HM007357.1, and HM007358.1) and JCV B (HM007353.1, HM007354.1, and HM007355.1), corresponding to the S, M, and L segments, respectively. Primers were mapped to each reference using corresponding BED files, which can be found at our JCVSeq protocols.io. Consensus genomes were generated using a threshold of 10 mapped reads per base and single-nucleotide variant frequency of 0.75^73^. We required 70% coverage across each of the three genomic segments for a sample to be included in phylogenetic analyses. All sequencing data can be found in the NCBI BioProject PRJNA1400525.

### Untargeted Metagenomic Sequencing

Following amplicon-based sequencing, we identified 35 samples with adequate (<32) CT values that fell below genome coverage thresholds for inclusion in our analysis. To ensure that these issues did not stem from sequencing primer mismatches, we additionally analyzed these samples using untargeted metagenomic sequencing methods as previously described^74^. We used 10 uL of the extracted mosquito pools as sample input and prepared all libraries using the Nextera XT preparation kit for Illumina. Prior to submitting, all libraries were checked for adequate concentration using the 1x dsDNA HS assay on the Qubit 4 (Thermo Fisher Scientific). Fragment sizes were measured using the High Sensitivity DNA Kit on the Bioanalyzer 2100 (Agilent). As in amplicon sequencing, libraries were submitted to the Yale Center for Genome Analysis for sequencing, at 5 million reads per sample on an Illumina NovaSeq 6000 (paired-end 150 bp). This yielded 7 additional samples, likely due to low viral concentration in mosquito pools. Sequencing data for these samples can be found in the NCBI BioProject PRJNA1400525.

### Alignment and Phylogenetic Analysis

The original 1961 JCV genomic sequence (GenBank accession numbers NC_043558.1, NC_043560.1, NC_043559.1 for the S, M, and L segments, respectively) has been previously identified as a reassortant virus^6,7^, so it could not be reliably used to root phylogenetic trees. Therefore, all divergence-based trees were rooted with the prototypic LaCrosse virus (LACV) sequence from 1960 (NC_077808.1, NC_077809.1, NC_077810.1) as an outgroup. Trees also included representative sequences of all viruses in the ICTV species classification *Orthobunyavirus jamestownense*, including Inkoo virus (INKV; KT288271.1, KT288270.1, KT288269.1), South River virus (SORV; KX817338.1, KX817337.1, KX817338.1), and Jerry Slough virus (JSV; KX817320.1, KX817319.1, KX817318.1). Additional published JCV sequences incorporated into the analysis included those from New Mexico (1977; MH900532.1, MH900531.1, MH900530.1), from Canada (1979; MH370820.1, MH370819.1, MH370818.1), and from Massachusetts (2016; MN135989.1, MN135990.1, MN135991.1). For all publicly available sequences, the medium segment accession number was used as a sample identifier across all segments to match tip names. All publicly available JCV sequences from Connecticut (from 1966, 2003, and 2004) were re-sequenced by our group as part of this project, and the sequences generated at Yale University were included in our analyses to avoid duplication.

When quantifying the number of publicly available sequences, we included JCV sequences with greater than 70% consensus coverage across each of the three genome segments. By this metric, 39 JCV genomes were publicly available. This number increases to 44 when considering full genomes taken from different timepoints of a fatal human infection. However, as this study primarily focused on the circulation of JCV in mosquitoes, these were excluded. The final alignments contained the 7 genomes available on GenBank, in addition to 32 sequences provided by the Wadsworth Center^22^. The other 658 sequences were generated at Yale using the amplicon-based sequencing method described above.

Initial phylogenetic analyses were performed using Nextstrain, an open source software that constructs maximum-likelihood phylogenetic trees and visualizes the evolutionary relationships between pathogen sequences^75^. The complete dataset includes 700 sequences. These trees were rooted with LACV and included the described INKV, SORV, and JSV sequences (3), JCV samples (7), provided samples from the NY Wadsworth Center (S segment: PX434836.1, PX439461.1-PX439491.1; M segment: PX441478.1-PX441509.1; L segment: PX445791.1-PX445822.1) (32), and novel JCV sequences generated at the Yale School of Public Health (658). Nextstrain builds for each segment of the JCV genome can be found at: https://github.com/grubaughlab/JCV-genomics.

To systematically designate JCV lineages, we used Autolin, an automated phylogenetic-based lineage assignment tool. Autolin assigns lineage labels using greedy maximization, integrating phylogenetic clustering and classification thresholds to designate lineages^27^. Nextstrain phylogenetic outputs for each JCV segment were uploaded to the Autolin web tool for lineage designation, with a minimum cluster size of 15. We further evaluated the generated lineage system for consistency across segments. While the system almost universally agreed across the medium and large segments, the small segment was significantly less stable. As such, after conducting several tree replicates, we decided that only the major lineage classification was appropriate for use on the small segment. More specific lineages can only be called across the medium and large segments. The resulting designation was further tested for robustness using other alignment tools. Multiple sequence alignments for each segment and concatenated genomes were generated using MAFFT v7.490^76^. Trees based on these alignments were built using IQ-TREE 2 (v2.3.6)^77^ with ultrafast bootstrap approximation using 1,000 replicates. We evaluated the resulting trees and their bootstrap values to ensure strong support (UFBOOT >95, SH-aLRT >80) for branches determining each lineage level ^77^. The bootstrap values supporting the A and B lineage nodes for the S segment were 100 and 98 respectively. For the M segment, the A lineage bootstrap values for minor lineages were as follows: 100 (A.1), 100 (A.1.1), 97 (A.1.2), and 100 (A.1.3). The support for M segment lineage B.1 was also 100. For the L segment, the A lineage bootstrap values for minor lineages were as follows: 100 (A.1), 99 (A.1.1), 100 (A.1.2), and 100 (A.1.3). The support for L segment lineage B.1 was also 100. All major lineages across each of the three segments, and minor lineages on the medium and large, were above this strong support threshold, allowing us to accept the lineage classification system as real. All trees and tanglegrams were visualized using “baltic”, an open source python package for parsing phylogenetic trees (https://github.com/evogytis/baltic).

### Identification of Reassortant Samples

To quantitatively assess the presence of reassortment events in our dataset, we employed Recombination Detection Program (RDP Beta 5.64)^78^. RDP uses a set of diverse algorithms for detecting recombination in nucleotide sequence alignments, applying both statistical and phylogenetic approaches (RDP, GENECONV, BootScan, MaxChi, Chimaera, SiScan, 3Seq)^78^. Although primarily used to identify recombination, RDP can also be leveraged to detect reassortment by analyzing incongruent signals across different segments in a concatenated genome^78^. To be considered a reassortment event, the reported breakpoints had to span a complete JCV genome segment (small, medium, or large). Only reassortment events supported by at least three independent methods with p-values below 0.05 were considered significant.

To further validate the reported reassortment events, we compared phylogenetic trees inferred by individual segments. These “tanglegrams” were generated using both Nextstrain and IQ-TREE 2 under the previously described conditions. To improve coherence across segments in the tanglegram, we implemented an “untangling” function that iteratively reorders internal nodes by aligning descendent tip positions across each tree. These visualizations confirmed the statistical outputs of RDP, demonstrating clear topological differences between samples across segments. Samples identified as reassortants via RDP had to be confirmed using the tree topology, and vice versa. All sequences identified as reassortments were pulled from the dataset before proceeding to Bayesian phylogenetic inference, as they would interfere with estimation of the evolutionary rate and other critical functions. A list of likely reassortants can be found in **Supplementary Table 3**. The RDP output for these samples and their associated statistical scores for each test can be found on the GitHub for this project at: https://github.com/grubaughlab/2025_paper_JCV.

### Temporal Signal Assessment

Initial tests of temporal signal were conducted on each segment using root-to-tip regression analyses conducted with the program TempEst v1.5.3^79^. However, we experienced significant difficulty observing a clear signal and calculating an overall evolutionary rate, even when separating segments by lineage. These issues were partially driven by the high level of genetic similarity between small segment genomes and long phylogenetic branch lengths. All identified reassortant sequences were excluded from analysis to eliminate possible interference with evolutionary rate estimation. We identified an unusually slow evolutionary rate across segments, with the S rate estimated as 5.74 x 10^-5^ substitutions per site per year (s/s/y), the M rate as 3.02 x 10^-5^ s/s/y, and the L rate as 2.634 x 10^-5^ s/s/y. To best capture temporal signal across the JCV genome, we chose to concatenate all three viral segments prior to analysis. To do so, all UTR regions were trimmed and the coding regions of all segments were strung together to create a single continuous sequence for each sample. Sequences were concatenated in S, M, and L segment order and re-aligned to ensure success. Due to the large divergence between lineages A and B, we chose to analyze the lineage trees separately. The lineage A alignment included all Northeast sequences in the largest monophyletic A.1 clade (468 sequences). The lineage B alignment included all Northeast sequences in the largest monophyletic B clade (124 sequences).

To assess the presence or absence of temporal signal and determine the best molecular clock to use for future analyses, we conducted a Bayesian Evolution of Temporal Signal (BETS) analysis for each lineage^80,81^. These, and all other phylogenetic analyses, were conducted using the software package BEAST X version 10.5.0, the high-performance phylogenetic likelihood library BEAGLE^82^, and the corresponding program BEAUti v10.5.0 to generate the XML input files^83^. To conduct a BETS analysis, we compared the phylogenetic outputs of our genomes with and without their associated sampling dates, in addition to testing the impacts of a strict and uncorrelated relaxed clock model^51^. In modeling the substitution rate, we implemented an HKY substitution model with gamma-distributed heterogeneity across 4 categories to accommodate for rate variation among sites, coupled with a constant population size coalescent prior^84–86^. We used all default priors as generated by BEAST X. Log marginal likelihood values were estimated through generalized stepping-stone (GSS) sampling to compare model performance^87^. The MCMC chain was set to run for 3 x 10^8^ steps, logging parameters every 1 x 10^4^ steps. The GSS was set for 3 x 10^7^ steps and checked regularly for convergence. Across lineages, we found that the uncorrelated relaxed clock model was the most appropriate for JCV time-scaling, and chose to use it for all subsequent analyses (**Supplementary Table 3** and **4)**. All analyses were assessed for convergence and estimated sampling size (ESS) values were evaluated in Tracer 1.7.3^88^. All XML and log files are available at: https://github.com/grubaughlab/2025_paper_JCV.

### Episodic evolutionary rate analyses

To infer potential episodic evolutionary rate dynamics, we employed epoch modeling within a Bayesian phylogeographic framework^89^. We implemented two configurations of the epoch model: (i) a model allowing distinct evolutionary rates for three periods: mid-May to end of October, November to mid-May (1997–2022), and a separate rate for the period prior to 1997; (ii) a similar model with alternating epochs defined as mid-May to mid-July and mid-July to mid-May (1997–2022). We specified CTMC conditional reference priors on the rates in the alternating epochs and a normal prior (mean=0.00003, SD=0.000005) for the ancestral rate. For this purpose, we extended the CTMC conditional reference prior for models involving more than one phylogeny simultaneously.

Specifically, we set the rate of the underlying gamma distribution to equal the epoch-specific average of the unknown tree lengths to retain the distributional tail-behavior derived across all trees^90^. The model is jointly fitted to independent phylogenies for lineages A and B, each using a flexible skygrid population size prior. Both evolutionary histories share an HKY substitution model with gamma-distributed rate variation.

### Discrete Trait Analyses

In order to interrogate JCV’s evolutionary history, we performed several discrete trait analyses on the A and B lineage trees. The first, reconstructing dispersal history in the Northeast US, modeled transitions between different states using an asymmetric forward-in-time CTMC model^91^. Each tip was labeled with its state of collection: CT (Connecticut), New York (NY), Massachusetts (MA), and Pennsylvania (PA). To identify well-supported transitions, we employed Bayesian stochastic search variable selection (BSSVS) to estimate non-zero transition rates between locations. We used all default priors as generated by BEAST X. MCMC analyses were run independently for each lineage for 1 × 10^9^ steps, with samples drawn every 1 × 10^5^ to characterize the posterior distribution. After analyses were assessed for adequate convergence, we retrieved and annotated maximum clade credibility (MCC) trees for visualization using TreeAnnotator v10.5.0^92^, keeping target node heights for all generated trees.

The second kind of discrete trait analyses were performed on Connecticut sequences, focused on transition rates between mosquitoes at the “generation-genus” level. The lineage A alignment included all lineage A.1.1 sequences in Connecticut, excluding reassortants (353 sequences). The lineage B alignment included all lineage B and B.0 sequences in Connecticut, excluding reassortants (120 sequences). Each tip was labeled according to the mosquito pool from which the virus was sampled, including “Uni-*Aedes*”, “Uni-*Coquillettidia*”, “Multi-*Aedes*”, “Multi-*Anopheles*”, and “Multi-Other”. The “Multi-Other” category captured three groups for which JCV was rarely detected, multivoltine *Culex*, *Culiseta*, and *Psorophora* species. Once again, we utilized a forward-in-time CTMC model and an uncorrelated relaxed molecular clock with an HKY substitution model^93^. We specified a Skygrid population model with 1-year grid intervals to flexibly capture temporal variation in effective population size over the surveillance period. We used all default priors as generated by BEAST X. MCMC analyses were run independently for each lineage for 1 × 10^9^ steps, with samples drawn every 1 × 10^5^ to characterize the posterior distribution. We additionally recorded Markov jumps^94^ for each mosquito group using the JumpHistoryLogger, once again logging every 1 × 10^5^ steps. We were also able to estimate the Markov Rewards, or time spent in each mosquito category, across the posterior distribution of trees^95^. After analyses were assessed for adequate convergence, we generated MCC trees for visualization using TreeAnnotator X^92^. All figures depicting Markov jumps were performed on the cumulative recorded jumps using the available Perl script on the BEAST website. In addition to the Markov jump analysis, we quantified the cumulative proportion of Markov rewards across the surveillance period (1997-2022) using the annotated MCC trees. Branch-specific reward annotations were parsed, divided in 0.1-year intervals, and aggregated by generation-genus mosquito group to estimate the reward proportions through time. We were then able to plot the temporal dynamics of these proportions using stacked area plots, restricted to the surveillance period. All analyses were assessed for convergence and ESS values with Tracer 1.7.3. All XML and log files are available at: https://github.com/grubaughlab/2025_paper_JCV.

### Surveillance and Seasonal Modeling

To assess the prevalence and seasonal patterns of JCV in Connecticut, we analyzed mosquito surveillance data collected by CAES from 1997 to 2022. This dataset includes arbovirus testing results, trapping metadata (location, date, and species), and mosquito abundance data. For analysis, data were filtered to include only the 26 mosquito species positive for JCV in Connecticut: *Aedes abserratus, Aedes aurifer, Aedes canadensis, Aedes cinereus, Aedes communis, Aedes excrucians, Aedes provocans, Aedes sticticus, Aedes stimulans, Aedes thibaulti, Aedes cantator, Aedes sollicitans, Aedes taeniorhynchus, Aedes triseriatus, Aedes trivittatus, Aedes vexans, Anopheles punctipennis, Anopheles quadrimaculatus, Anopheles walkeri, Psorophora ferox, Coquillettidia perturbans, Culex erraticus, Culex salinarius, Culex restuans, Culiseta melanura*, and *Culiseta morsitans*^6,7^. Mosquito abundance was calculated as the number of individuals per trap per night, grouped by date, site, trap type, and species. JCV positivity was encoded as a binary variable, with positive pools assigned a value of 1 and negative pools 0. The pooled infection rate (pIR), representing the estimated proportion of JCV-infected mosquitoes, was calculated using maximum-likelihood estimation (MLE) via the CDC PooledInfRate package (https://github.com/CDCgov/PooledInfRate). This method accounts for variability in the number of positive mosquitoes per pool, estimating the infection probability per epi-week. To quantify JCV’s transmission potential, we calculated the vector index (VI) as the product of pIR and mosquito abundance. VI provides an estimate of the relative pathogen load in different mosquito groups. pIR and VI were both calculated at multiple taxonomic levels, including species, genus-generation categories, and overall generation type. Long-term (annual) and short-term (weekly) trends in JCV dynamics were also analyzed to assess seasonal fluctuations and inter-annual variation in virus prevalence. Code used in this analysis can be found at: https://github.com/grubaughlab/2025_paper_JCV.

### Continuous Phylogeographic Reconstruction

To map the movements of JCV over time and space, we implemented a continuous phylogeographic analysis for each lineage, A and B. Each sequence was assigned a unique latitude and longitude value. For sequences originating from the same trapping location, we generated random latitude and longitude from non-overlapping areas of uncertainty surrounding each trap. These areas had a circular radius of 650 meters. In order to account for uncertainty in the phylogenetic tree, while also limiting the required computation power for the analyses, we used 1000 empirical trees from the previous discrete trait analyses (A and B). Using these, we ran a single continuous phylogeographic analysis that included both lineage A and B. Both tree models were estimated jointly with a single MCMC, while each lineage retained its own empirical tree distribution, multivariate continuous diffusion model, and branch-specific diffusion rates. The MCMC analysis was run for 1 x 10^8^ steps, with operator weights assigned to tree distribution moves and Hamiltonian Monte Carlo updates of diffusion rate parameters^96^. Logging was performed at multiple levels. We sampled the phylogenetic trees every 1 x 10^3^ steps. Separate log files were maintained for lineages A and B to allow lineage-specific reconstruction of spatial dynamics. All analyses were assessed for convergence and ESS values with Tracer 1.7.3. All XML and log files are available at: https://github.com/grubaughlab/2025_paper_JCV.

We used the R package “seraphim”^97^ to visualize the continuous phylogeography and estimate dispersal statistics for each lineage. For each lineage (A and B) and sampled posterior tree, we estimated the weighted diffusion coefficient (km^2^/year) by deriving branch-specific movement distances and times from spatially annotated phylogenetic nodes^98^. We also estimated an isolation-by-distance signal computed as the Pearson r_P_ correlation between patristic and log-transformed geographic distances between phylogenetic tips^50^. We additionally repeated the analyses focusing exclusively on the surveillance period in Connecticut, from 1997 to the present, to capture phenomena specific to this time. Dispersal statistics across each of these analyses can be found in **Supplementary Table 5**.

### Landscape Phylogeographic Analyses

To further analyze the impact of environmental factors on the dispersal history of JCV, we conducted two different landscape phylogeographic analyses. These methods, performed using the R package “seraphim”, tested the impact of several variables on the dispersal location^99^ and diffusion velocity^100^ of inferred viral lineages. Both of these analyses rely on the comparison between spatially annotated trees inferred by the continuous phylogeographic analysis and the corresponding spatially annotated trees generated under a null dispersal model. Under this null dispersal model, in which no environmental factor was associated with the dispersal dynamic of viral lineages, the phylogenetic branches of each inferred tree are randomized within the study area while preserving the tree topology, maintaining the inferred position of the most ancestral node, and preventing nodes from falling into non-accessible (i.e., sea/water) areas^100,101^. This randomization procedure thus provides a baseline diffusion process, which we can then compare with the patterns inferred by our continuous phylogeography, investigating the influence of different environmental factors formalized as rasters (i.e. geo-referenced grids of environmental values) on the dispersal dynamic of JCV lineages. We retrieved rasters for the following environmental variables potentially biologically relevant to the ecology of JCV: elevation (Shuttle Radar Topography Mission), forest areas, croplands, urban areas, and water areas (International Geosphere Biosphere Programme), annual temperature, and annual precipitation across the state of Connecticut (WorldClim database)^52^.

First, we analyzed how our listed environmental variables were associated with the dispersal location of JCV lineages, testing whether inferred viral lineages tended to preferentially circulate or avoid circulating in specific environmental conditions. For each environmental variable, we generated a posterior distribution of the mean values extracted at all phylogenetic node positions. These mean values could then be compared across the inferred and randomized trees, computing a Bayes factor (BF) support associated with each variable using the following formula: BF = (*p_e_* /(1−*p_e_*))/(0.5/(1−0.5), where *p_e_* represents the frequency at which inferred values exceeded or fell below randomized values. We interpreted a BF >20 as strong statistical support^102^. The results of this analysis can be found in **Supplementary Table 6**.

We then investigated to what extent these environmental variables could be either negatively or positively associated with some heterogeneity in the inferred diffusion velocity of JCV lineages; testing each environmental factor once as a potential resistance (decreasing diffusion velocity) and as once as a potential conductance (increasing diffusion velocity) factor, respectively. For each tree branch across inferred and randomized trees, we calculated environmental distances using both the least-cost^103^ and Circuitscape^104^ path models. For this, the original raster cell values were preliminary transformed using the following formula: *v_t_*= 1+*k*(*v_0_*/*v_max_*), where *v_t_*and *v_0_* correspond to the transformed and original cell values, respectively, *v_max_* to the maximum cell value recorded in the raster, and *k* to a rescaling parameter. We considered three distinct values for *k* (*k* = 10, 100, and 1000), which allowed testing different strengths of conductance or resistance relative to a null raster where all accessible cells have a value equal to “1”. For each resulting environmental raster, we estimated the *Q* statistic measuring the correlation between branch durations and the associated squared environmental distances^100^. An environmental raster was considered a potential explanatory variable when its associated posterior distribution of *Q* values was considered as positive, i.e. when 90% of the *Q* values were >90%^105^. In such cases, the statistical support for the *Q* distribution was formalized as a BF support by comparing the posterior distribution of *Q* values with its corresponding distribution obtained when computing environmental distances for the randomized trees^106^. The results of this analysis can be found in **Supplementary Table 7**.

### General Additive Modeling

In addition to our landscape analyses, we employed a modeling-based approach to test the impact of statewide ecological factors on virus diffusion over time. To do so, we fitted Bayesian generalized additive models (GAMs) to the posterior weighted diffusion coefficient (WDC) estimated for each lineage. We fitted two separate models for lineages A and B, considering both a 1-year and 5-year sliding window, yielding 2,000 posterior data frames per lineage (4,000 in total). Each data frame contained WDC estimates post-1997, the number of phylogenetic branches corresponding to each estimate, and the year as set by the sliding window. We joined these outputs to statewide ecological covariates in Connecticut, including mean, maximum, and minimum temperature, precipitation, as well as the z-index (a measure of anomalies in soil moisture). Mosquito-related variables included the total abundance of each generation-genus category (univoltine *Aedes* and *Coquillettidia*, multivoltine *Aedes*, *Anopheles*, and others) as surveyed by CAES since 1997. In addition, we included lineage-specific counts of JCV-positives by mosquito group, based on the lineage calls and pool identity of all our positive samples per year. These were tested for covariation prior to model input to avoid autocorrelation between ecologically linked variables. We chose to proceed with measures of maximum temperature, precipitation, and our JCV-positive mosquito abundance. To account for uncertainty in tree topology and diffusion estimated, the number of phylogenetic branches was included as a smoothing spline (s(number_of_branches, k = 5, bs = “tp”)). The sliding window parameter (1 or 5) was also included as a random intercept to consider for any variation between window sizes. All models were implemented with the R package “brms” (v2.21.0) using Hamiltonian Monte Carlo Sampling. The final model was run with four chains of 2,400 iterations each, using 1,200 of these for warmup. Convergence was assessed using both the potential scale reduction factor (R < 1.01) and effective sampling size (Bulk ESS and Tail ESS > 1,000 for each variable).

Posterior means and 95% HPD intervals were summarized for all parameters and visualized. Code used in this analysis can be found at: https://github.com/grubaughlab/2025_paper_JCV.

## Data Availability

All sequencing data can be found in the NCBI BioProject PRJNA1400525 and the Supplementary Tables.

## Code Availability

The code and phylogenetic files (XML, log, and RDP files) can be found at: https://github.com/grubaughlab/2025_paper_JCV.

## Acknowledgements

We acknowledge the county health departments in NYS and the NYS Bureau of Communicable Disease Control for mosquito collections, E. Holmes for feedback on the evolutionary rates, and S. Taylor and P. Jack for technical advice. This publication was made possible by the National Institute of Allergy and Infectious Diseases of the National Institutes of Health (NIH) under Award Number DP2AI176740 (NDG), *Fonds National de la Recherche Scientifique* (F.R.S.-FNRS, Belgium) (SD), Research Foundation - Flanders (“Fonds voor Wetenschappelijk Onderzoek - Vlaanderen,” G0E1420N, G098321N) (GB), European Union Horizon 2023 RIA project LEAPS (grant agreement no. 101094685) (GB), DURABLE EU4Health project 02/2023-01/2027 which is co-funded by the European Union (call EU4H-2021-PJ4) under Grant Agreement No. 101102733 (GB), National Science Foundation award number DBI 2515340 (CJC), and National Institute of General Medical Sciences of the NIH under Award Number 1S10OD030363-01A1 awarded to the Yale Center for Genome Analysis. The findings and conclusions in this report are those of the author(s) and do not necessarily represent the official position of the NIH.

## Author Contributions

E.B, A.T.C., C.B.F.V., V.H., P.M.A., and N.D.G. conceived and designed the study. A.B.B., M.J.M., T.A.P., J.J.S., T.G.A., J.F.A., K.A.N., J.G.M., A.P.D., S.M.R., G.X., G.S., K.J.P., M.L.M., A.T.C., and P.M.A. collected and provided the data and samples. E.B., N.M.F., M.I.B., and C.B.F.V. designed and conducted the sequencing. E.B., S.D., P.L., M.A.S., R.L., F.G., C.J.C., G.B., V.H., and N.D.G. conducted the analyses and interpreted the data. E.B. led the writing with N.D.G. and all authors reviewed, edited, and approved the final version.

## Extended Data Figures and Supplementary Tables

**Extended Data Fig. 1.**
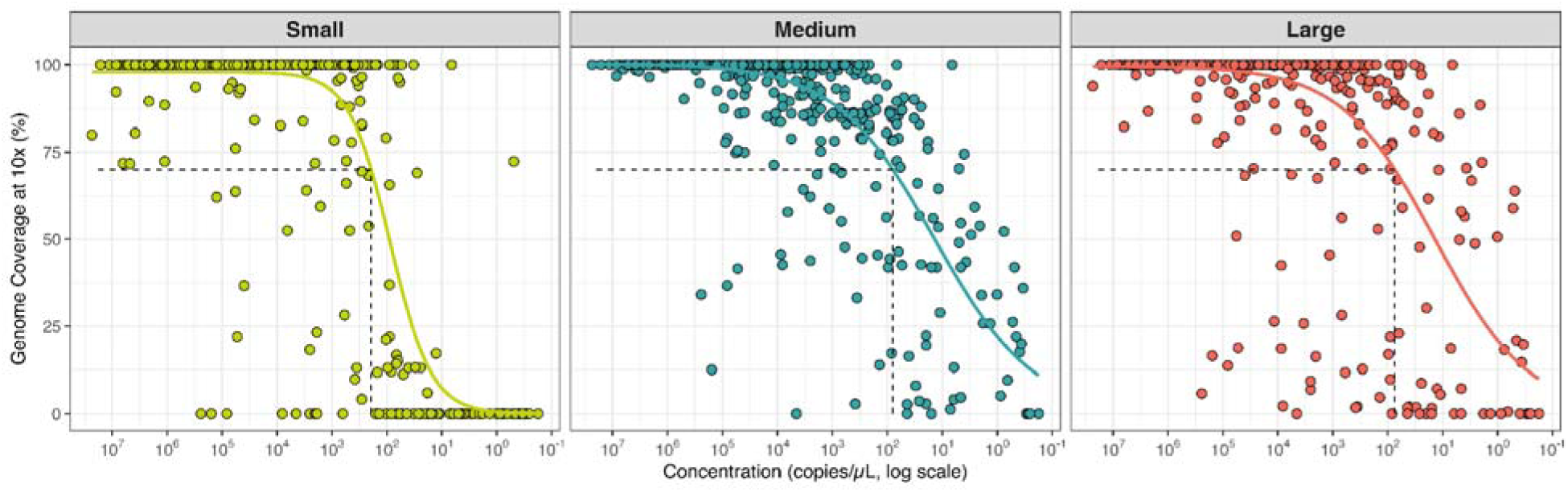
JCV sequencing validation. Genome coverage at 10x depth is plotted against input viral RNA concentration (copies/μL) as reported by qPCR for each of the genomic segments: small, medium, and large. Each point represents a sample extracted from mosquito pools in the United States. The x-axis is presented on a lo scale, with higher concentrated samples on the left and lower concentrations on the right (descending concentration scale). Horizontal lines indicate the 70% threshold for genome coverage, the metric used for inclusion in this study. Vertical dashed lines indicate the estimated detection limit for each segment, calculated using a sigmoidal fit of coverage vs. concentration, representing the lowest concentration at which predicted coverage reaches 70%. Thes calculated values were 193 copies/μL, 79 copies/μL, and 74 copies/μL for the small, medium, and large segments respectively.

**Extended Data Fig. 2.**
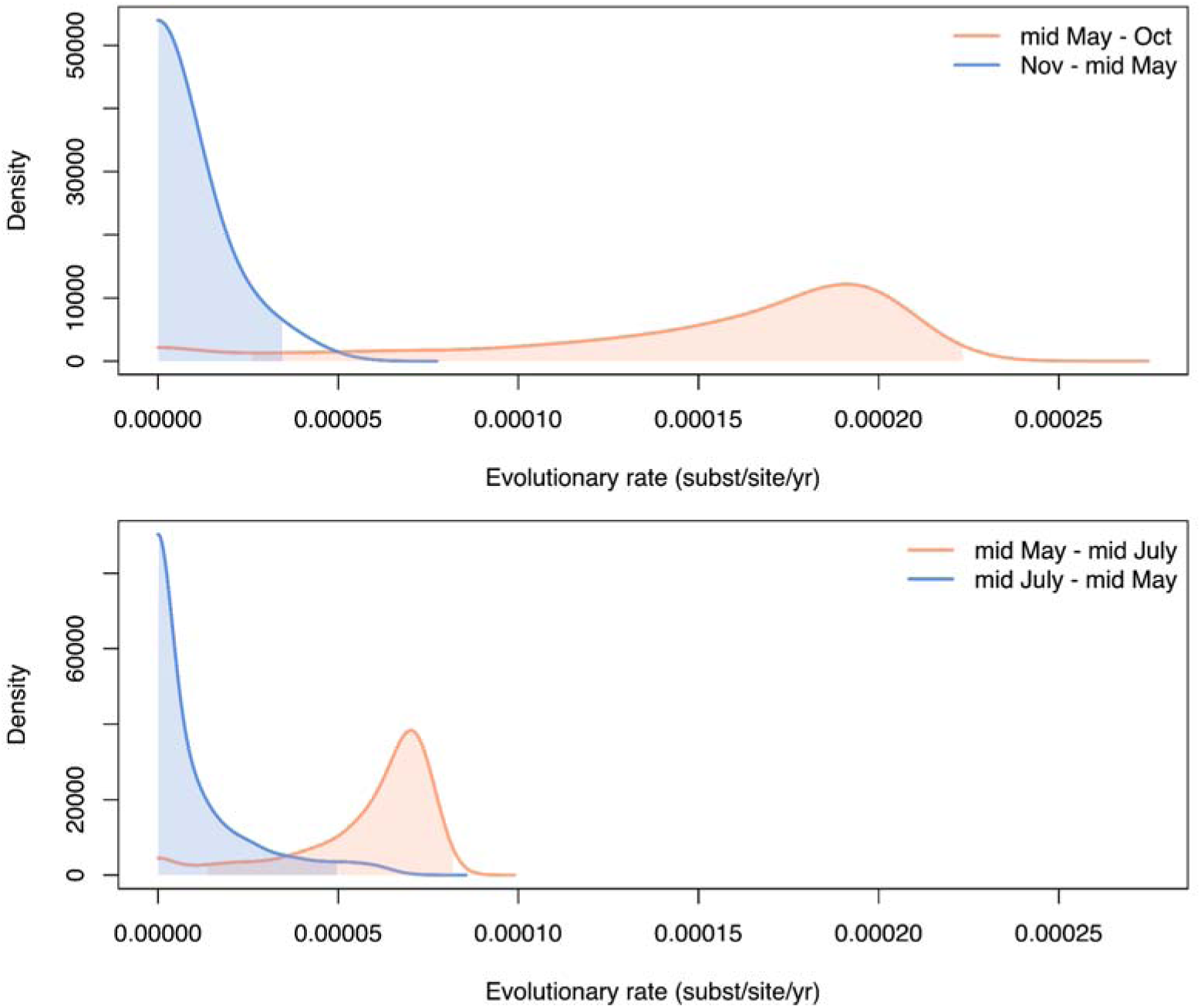
Posterior distributions of JCV evolutionary rate estimates under alternative seasonal epoch models. Posterior density distributions of substitution rate estimates for JCV inferred using two epoch-model configurations jointly fitted to lineages A and B. Top panel: Epochs defined as Mid-May to the end of October and November to mid-May (1997-2022), with a separate ancestral rate estimated for the period prior to 1997. Bottom panel: Epochs defined as mid-May to mid-July and mid-July to mid-May (1997-2022). Rates were estimated using Bayesian epoch modeling with CTMC conditional reference priors on epoch-specific rates and a normally distributed prior on the ancestral rate. Density is shown on the y-axis and evolutionary rate (substitutions per site per year) o the x-axis.

**Extended Data Fig. 3.**
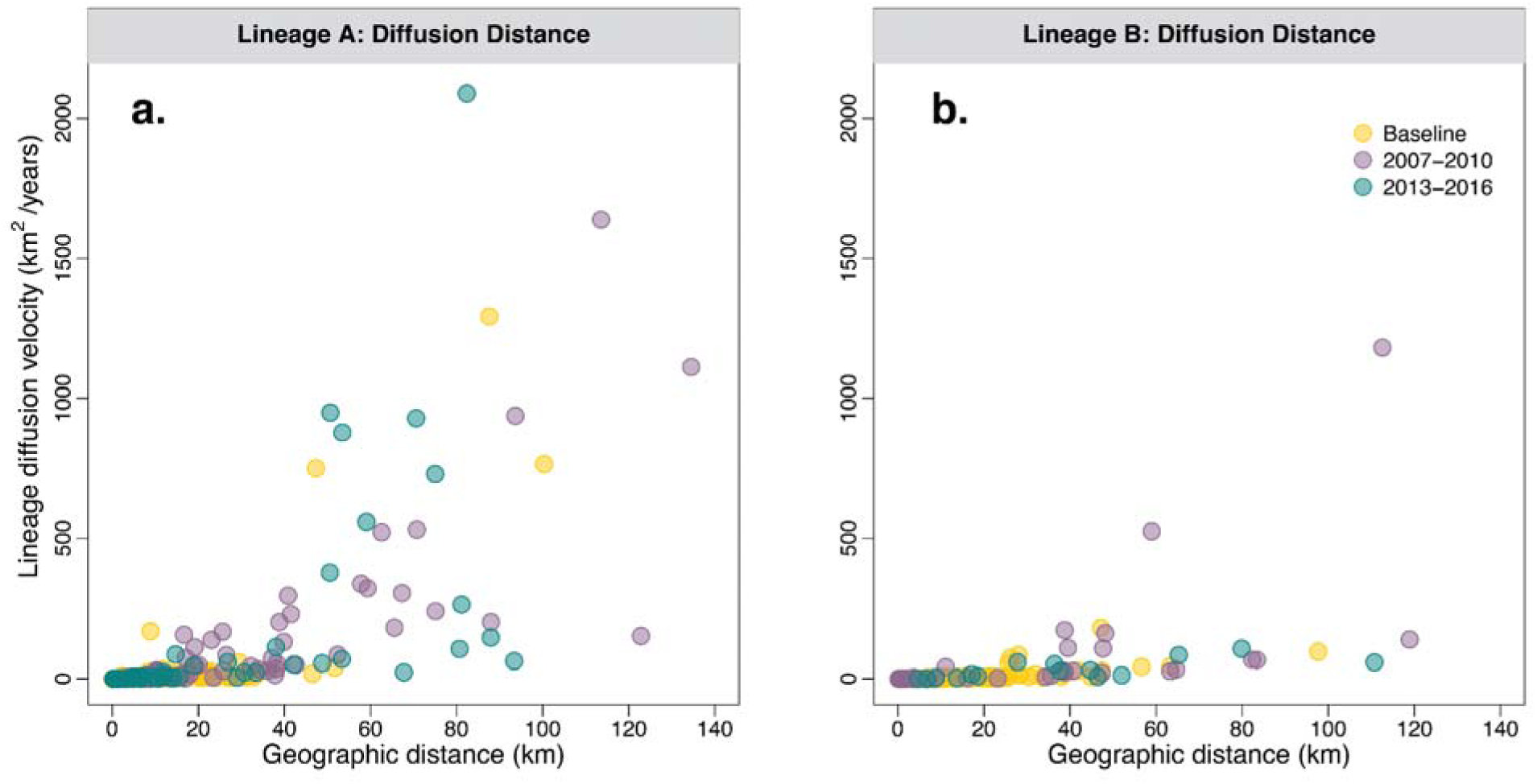
Dispersal distance versus lineage diffusion for lineages A and B. Plots depict the relationship between geographic distance (km) and lineage diffusion (km2/year), assessing whether peaks in the weighted diffusion coefficient (WDC) is driven by rapid, long-distance dispersal events. Each point represents a branch from the maximum clade credibility (MCC) tree for lineage A (**a**) and lineage B (**b**). Points are colored by temporal windows: baseline (pre-2007, yellow), 2007-2010 (purple), and 2013-2016 (teal, lineage A only). Results indicate that peaks in WDC are driven by a limited number of rapid, long distance dispersal events.

**Extended Data Fig. 4.**
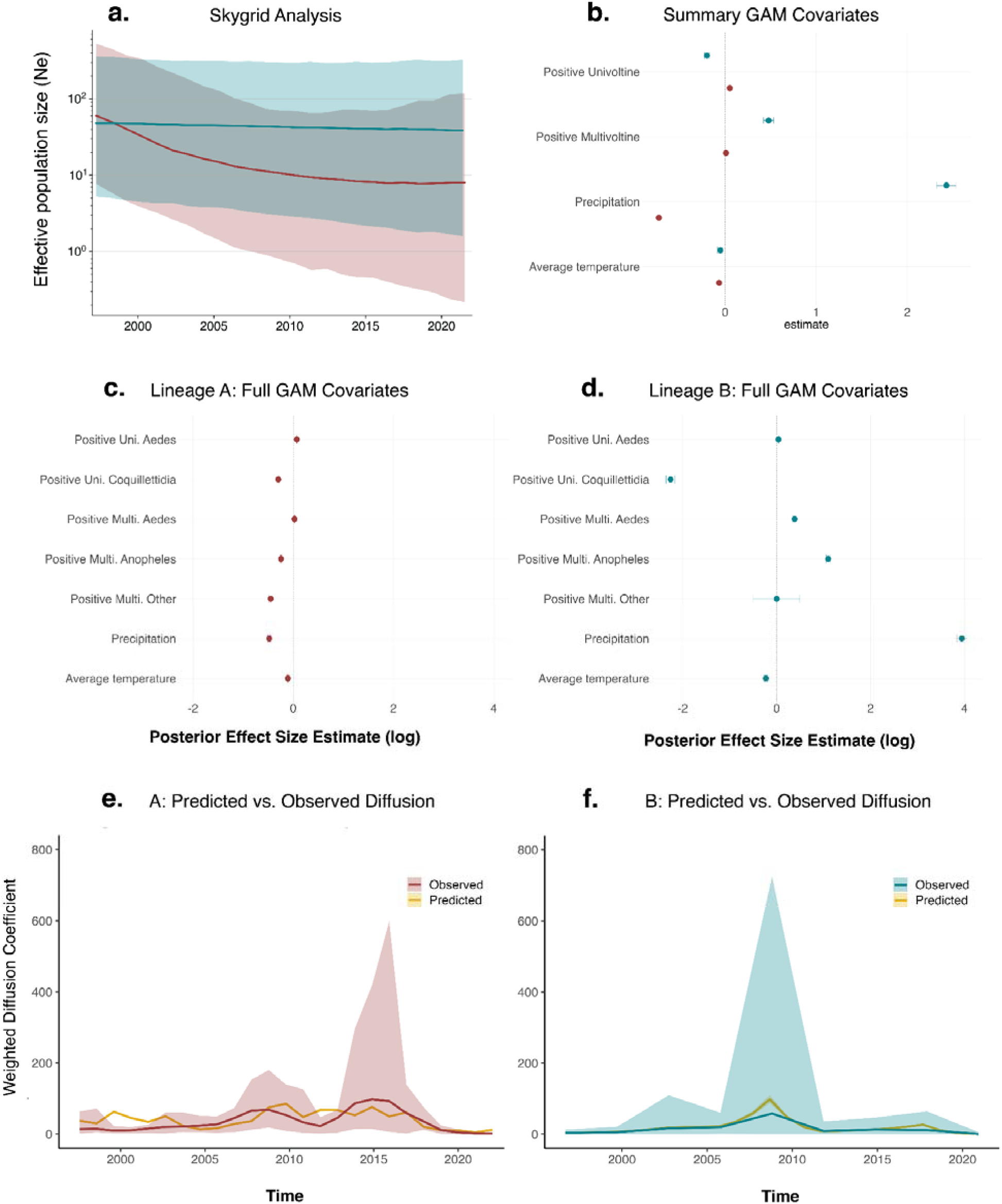
Demographic history and ecological drivers of JCV spatial diffusion. (**a**) Skygrid reconstruction of the evolution of the overall effective size of viral lineages A and B over the surveillance period (1997-2022); (**b**) Estimated log effect size of ecological variables associated with JCV WDC per year, using aggregated univoltine and multivoltine mosquito categories; (**c**) Estimated log effect size of ecological variables associated with lineage A WDC per year, by mosquito generation and genus. Branch number coefficient (not shown) estimated at 3.42 (3.04-3.83); (**d**) Estimated log effect size of ecological variables associated with lineage B WDC per year, by mosquito generation and genus. Branch number coefficient (not shown) estimated at 0.38 (0.34-42); (**e**) Predicted lineage A WDC (km^2^/year) as generated by Bayesian GAM vs. observed WDC; (**f**) Predicted lineage B WDC (km^2^/year) as generated by Bayesian GAM vs. observed WDC.

