## Supplementary Tables for "Evolutionary history of Jamestown Canyon virus disentangles complex multi-vector ecology"

**Supplementary Table 1.** **JCVSeq sequencing primers.** All primers with names starting with “NYS” represent UTR segment specific primers provided by the New York State Department of Health’s Wadsworth Center. All primers with names starting with “Yale” were newly developed as a part of this study using PrimalScheme. Each primer is labeled with the targeted segment, nucleotide position on the reference sequences, and the major lineage A or B. Primer pairs are overlapping, with corresponding forward and reverse pairs labeled with matching numbers.

| **Primer Name** | **Sequence** |
| --- | --- |
| NYS_JCVSMF1 | AGTGTACTACCAAGTATAGAAAACGTTCA |
| NYS_JCVSM991R1 | AGTAGTGTGCTCCACTGAATACATTTAA |
| NYS_JCVMF1 | AGTAGTGTACTACCAAGTATAGAAAACGTT |
| Yale_JCVM837R1B | GGGTGATAAACTAACCCGCACA |
| Yale_JCVM837R1A | GGGTGATACACCAAACCACATAGT |
| Yale_JCVM582F2B | ACCAACATATGTCATGTGTACGGT |
| Yale_JCVM582F2A | ATCAGCATATGTCATGCGTACGAT |
| Yale_JCVM1367R2B | TGTGTACTATCAATCCAGTAACATCTTCA |
| Yale_JCVM1367R2A | TATGAACCATTAACCCAGTAATGTCTTCA |
| Yale_JCVM1257F3B | GTGATATGTATCATGAAAAAGCCGGT |
| Yale_JCVM1257F3A | GTGACATGTACCATGAAAAAGCTGG |
| Yale_JCVM2057R3B | TGTCTTTTCGGGCCATAGCTTC |
| Yale_JCVM2057R3A | TATCTTTGCGAGCCATAGCTTCTG |
| Yale_JCVM1910F4B | TGGGATTTTGCAAATGAAATGAAGACA |
| Yale_JCVM1910F4A | TGGGATTTCGCAAATGAAATGAAAACT |
| Yale_JCVM2747R4B | TACTGCATCTGGGACTAAGGCA |
| Yale_JCVM2747R4A | TGTTGCACCTAGGGCTTAGACA |
| Yale_JCVM2611F5B | TGGGAACTGTGATGTTCAAGAAAATG |
| Yale_JCVM2611F5A | TGGAAATTGCAATGTTCAAGAAAATGATT |
| Yale_JCVM3440R5B | TGGGAGTTATAGCATTTAAGTTTGTGC |
| Yale_JCVM3439R5A | AGGAGTTACAGCATTTAAATTAGTGCAGTA |
| Yale_JCVM3303F6B | TCTTCGGGTCCTGTCAAGACATTA |
| Yale_JCVM3299F6A | TGTGTATTTGGATCCTGTCAAGATATCATA |
| Yale_JCVM4134R6B | TGGTCTACTGGTGCAAGCTCTAG |
| Yale_JCVM4134R6A | TGGTCAACTGGTGCAAGTTCTAA |
| Yale_JCVM3680F7B | GACAACGATTACCAAGCTTGCAA |
| Yale_JCVM3680F7A | GACAATGATTATCAAGCTTGCAAATTTCT |
| NYS_JCVM4510R2 | AGTAGTGTGCTACCAAGTATATCTAAATGA |
| NYS_JCVLF1 | AGTAGTGTACTCCTATTTACAAAACTTACAAATAC |
| Yale_JCVL873R1B | GGGCTGTTTGTAATCTCCAGTGG |
| Yale_JCVL873R1A | GGGCTGTTTGTAGTTCCCTGT |
| Yale_JCVL746F2B | CATATGAATCAGAAAGATGGAACACTAACC |
| Yale_JCVL746F2A | CATACGAATCAGAGAGGTGGAACA |
| Yale_JCVL1577R2B | CATTGCCAGAATTTGGTCTCAAAAATAG |
| Yale_JCVL1577R2A | CACTGCCAGAACCTAGTCTCAAA |
| Yale_JCVL1177F3 | AGCTGGCGACAAGTTATGAATAAGA |
| Yale_JCVL2014R3B | GTGCAGGCTCAGTTAGAGATAACA |
| Yale_JCVL2014R3A | GTGCAGGCTCAGTTAAGGACAA |
| Yale_JCVL1680F4B | TGCATTAGTTTACCCTTCAGCTGAT |
| Yale_JCVL1680F4A | TGCACTAGTCTACCCATCTGCA |
| Yale_JCVL2453R4B | AGGATTTGTAAATTAACTGTTTGCTTGGT |
| Yale_JCVL2453R4A | AGGATCTGCAAATTGACAGTTTGTTT |
| Yale_JCVL2315F5B | CCAAAGGTCTGCATGAGAAGCA |
| Yale_JCVL2315F5A | CTAAGGGTTTACATGAGAAACATCATGTT |
| Yale_JCVL3121R5B | CCCTATTTTTTTGCCTAGTAGTTTCAACT |
| Yale_JCVL3121R5A | CCCTATTCTTTTGTCTTGTAGTCTCAAC |
| Yale_JCVL2940F6B | TGTAGGAGAATATGAAGCCAAAATGTGC |
| Yale_JCVL2940F6A | TGTGGGAGAATACGAGGCTAAAATG |
| Yale_JCVL3744R6 | TGTCTTTTTCATATTGGCTTGACATCC |
| Yale_JCVL3638F7B | CAATAGTTCAAGATAAGGCCCCTGA |
| Yale_JCVL3638F7A | CGATAGTTCAAGACAAGGCTCCA |
| Yale_JCVL4432R7B | CATTCATGCCTCCTGGTGATGATA |
| Yale_JCVL4432R7A | CATTCATTCCTCCTGGTGATGACA |
| Yale_JCVL4301F8B | TGGGTGAGACAAGTGATATGAGGG |
| Yale_JCVL4301F8A | TGGGTGAAACAAGCGATATGAGAG |
| Yale_JCVL5127R8B | TCTTGTCATTTCTTTCAATTCAAACACAAT |
| Yale_JCVL5126R8A | CTCGTCATCTCCTTCAGTTCAAAAAC |
| Yale_JCVL4975F9B | GCAGACCCAACAGAGATGTCAA |
| Yale_JCVL4975F9A | GCTGATCCAACAGAGATGTCAAGA |
| Yale_JCVL5804R9B | GCCATATTTTCGAATTTTAGACCATGC |
| Yale_JCVL5804R9A | GCCATATTTTCAAATTTCAAGCCATGT |
| Yale_JCVL5681F10B | TTGGTGAAGACAACAAGCTAACTTATTC |
| Yale_JCVL5681F10A | TCGGTGAAGACAATAAGCTAACCTAC |
| Yale_JCVL6499R10B | TAGCCTCTAAGCAACCTAAGTTTTCA |
| Yale_JCVL6499R10A | TGGCTTCTAAACAACCTAAATTTTCTGG |
| Yale_JCVL6087F11B | AAAGAAAGCACATTTTAGCAAAATGGTATC |
| Yale_JCVL6087F11A | AAAGAAGGCGCATTTCAGCAAAA |
| NYS_JCVL6957R2 | AGTGTGCTCCTATTTACAAATATATACTATAAGC |

**Supplementary Table 2.** **Reassortant JCV samples as identified by the Recombination Detection Program (RDP5), 1960-2023.** Each row represents a sequence flagged as a possible reassortant. The “Small”, “Medium”, and “Large” columns indicate the lineage assignment for each of the three genomic segments (S, M, and L) based on segment-specific phylogenetic analyses.

| **Sample** | **Sampling date** | **Sampling location** | **Mosquito species** | **Small**  **lineage** | **Medium lineage** | **Large lineage** |
| --- | --- | --- | --- | --- | --- | --- |
| KX817316 | 1961 | Jamestown, Colorado | *Culiseta inornata* | B-related | B-related | B-related |
| KX817319 | 1963 | Kern, California | *Culiseta inornata* | B-related | B-related | B-related |
| MH900531 | 1977 | Unknown, New Mexico | *Culiseta inornata* | B-related | B-related | B-related |
| 117 | 2003-07-22 | Windham, Connecticut | *Aedes canadensis* | A | A.2 | A.2 |
| 221 | 2005-08-08 | Cass, North Dakota | *Aedes dorsalis* | A | A.2 | A.2 |
| 225 | 2005-08-09 | Williams, North Dakota | *Aedes vexans* | A-related | A-related | A-related |
| 226 | 2005-08-09 | Williams, North Dakota | *Aedes melanimon* | A-related | A.1 | A-related |
| 343 | 2009-07-01 | Stonington, Connecticut | *Aedes aurifer* | A | A.1.1 | B.1 |
| 344 | 2009-07-01 | Stonington, Connecticut | *Aedes cantator* | A | B.1 | A.1.1 |
| JC13 | 2009 | Erie, New York | *Aedes stimulans* | A | A.2 | A.1.2 |
| 688 | 2011-06-30 | Erie, New York | *Aedes canadensis* | A | A.1.2 | A.1.2 |
| JC333 | 2022 | Onondaga, New York | *Coquillettidia perturbans* | A | A.1.3 | A.1.2 |

**Supplementary Table 3. Lineage A Bayesian Evaluation of Temporal Signal.** GSS (Generalized Stepping-Stone) values represent the log-marginal likelihoods for models without (no sampling dates) and with (sampling dates) temporal information. Positive GSS values indicate strong support for the inclusion of sampling dates, reflecting the presence of temporal signal in the dataset.

|  | **Strict Clock** | **Relaxed Clock** |
| --- | --- | --- |
| **No sampling dates** | -63,497.33 | -63,475.43 |
| **Sampling dates** | -59,868.39 | -59,867.17 |
|  | **3,628.94** | **3,608.26** |

**Supplementary Table 4. Lineage B Bayesian Evaluation of Temporal Signal.** GSS (Generalized Stepping-Stone) values represent the log-marginal likelihoods for models without (no sampling dates) and with (sampling dates) temporal information. Positive GSS values indicate strong support for the inclusion of sampling dates, reflecting the presence of temporal signal in the dataset.

|  | **Strict Clock** | **Relaxed Clock** |
| --- | --- | --- |
| **No sampling dates** | -29,477.73 | -29,460.53 |
| **Sampling dates** | -29,443.89 | -29,446.19 |
|  | **33.84** | **14.34** |

**Supplementary Table 5. Dispersal statistics of JCV lineages in Connecticut.** Key spatial diffusion metrics were estimated from continuous phylogeography analyses, calculated using seraphim. For each metric, we here report the median value along with the corresponding 95% highest posterior density (HPD) interval in brackets. Values were aggregated over 1000 posterior trees and calculated for both major lineages. Here, the full surveillance time describes the total duration of spread from the root to 2022. The surveillance period is defined as 1997-2022, as determined by Connecticut Agricultural Experiment Station (CAES) sampling.

| **Lineage** | **Time period** | **Weighted diffusion coefficient (km^2^/year)** | **Isolation-by-distance signal** |
| --- | --- | --- | --- |
| A | Full | 32.20 [27.73, 37.47] | 0.19 [0.18, 0.21] |
| A | Surveillance | 57.22 [50.03, 64.17] | 0.01 [-0.02, 0.05] |
| B | Full | 19.33 [13.25, 26.99] | 0.08 [0.07, 0.10] |
| B | Surveillance | 62.38 [49.12, 78.31] | 0.02 [-0.04, 0.09] |

**Supplementary Table 6: testing the association between the dispersal locations of JCV lineages and a set of environmental factors.** We report Bayes factor (BF) support for the association between environmental values and tree node locations. The results are based on 1,000 posterior trees obtained by spatially explicit phylogeographic inference. Following Kass & Raftery (1995), we consider a BF value >20 as strong support (in bold).

| **Environmental factor** | **Tendency of viral lineages to**  **avoid circulating within specific environmental conditions** | **Tendency of viral lineages to preferentially circulate within specific environmental conditions** |
| --- | --- | --- |
| Elevation | **24.0** | 0.0 |
| Forest areas | 10.6 | 0.1 |
| Croplands | **99.0** | 0.0 |
| Urban areas | 0.1 | 12.7 |
| Annual mean Temperature | 0.2 | 3.5 |
| Annual precipitation | 0.1 | 7.3 |

**Supplementary Table 7:** testing the association between the diffusion velocity of JCV lineages and a set of environmental factors. The results are based on 1,000 posterior trees obtained by spatially explicit phylogeographic inference when considering the least-cost path model, but only on 100 posterior trees when considering the Circuitscape path model. “C” and “R” indicate if the considered environmental raster was considered as a conductance (“C”) or resistance factor (“R”), and *k* is the rescaling parameter used to transform the initial raster (see the text for further details). For regression coefficients and *Q* values we report both the median estimate and the 95% HPD interval. The Bayes factor (BF) supports are only reported when *p*(*Q* > 0) is at least 90%. Following Kass & Raftery (1995), we consider a BF value >20 as strong support (none in this case).

| **Path model** | **Environmental** **factor** | ***k*** | **Regression coefficient** | ***Q* statistic** | ***p*(*Q* > 0)** | **BF** |
| --- | --- | --- | --- | --- | --- | --- |
| **Least** | elevation (C) | 10 | 0.021 [0.008, 0.046] | -0.004 [-0.013, 0.002] | 0.113 | - |
| **-cost** |  | 100 | 0.016 [0.005, 0.039] | -0.009 [-0.023, 0.003] | 0.076 | - |
| **algorithm** |  | 1000 | 0.013 [0.003, 0.035] | -0.011 [-0.029, 0.003] | 0.047 | - |
|  | elevation (R) | 10 | 0.025 [0.010, 0.052] | 0.001 [-0.012, 0.012] | 0.553 | - |
|  |  | 100 | 0.016 [0.006, 0.035] | -0.007 [-0.031, 0.007] | 0.177 | - |
|  |  | 1000 | 0.014 [0.005, 0.031] | -0.009 [-0.033, 0.006] | 0.129 | - |
|  | forest areas (C) | 10 | 0.024 [0.003, 0.054] | -0.001 [-0.025, 0.013] | 0.464 | - |
|  |  | 100 | 0.017 [0.001, 0.046] | -0.006 [-0.030, 0.012] | 0.204 | - |
|  |  | 1000 | 0.016 [0.000, 0.044] | -0.008 [-0.031, 0.012] | 0.167 | - |
|  | forest areas (R) | 10 | 0.016 [0.006, 0.038] | -0.007 [-0.029, 0.005] | 0.121 | - |
|  |  | 100 | 0.014 [0.004, 0.032] | -0.010 [-0.034, 0.003] | 0.061 | - |
|  |  | 1000 | 0.013 [0.004, 0.032] | -0.011 [-0.034, 0.002] | 0.055 | - |
|  | croplands (C) | 10 | 0.036 [0.015, 0.069] | 0.011 [-0.001, 0.027] | 0.963 | 10.6 |
|  |  | 100 | 0.045 [0.019, 0.093] | 0.020 [-0.009, 0.059] | 0.915 | 8.3 |
|  |  | 1000 | 0.040 [0.013, 0.081] | 0.014 [-0.019, 0.057] | 0.816 | - |
|  | croplands (R) | 10 | 0.025 [0.001, 0.050] | 0.000 [-0.006, 0.007] | 0.521 | - |
|  |  | 100 | 0.006 [0.000, 0.025] | -0.018 [-0.04, -0.002] | 0.019 | - |
|  |  | 1000 | 0.001 [0.000, 0.009] | -0.023 [-0.049, -0.007] | 0.000 | - |
|  | urban areas (C) | 10 | 0.012 [0.003, 0.031] | -0.012 [-0.035, 0.000] | 0.028 | - |
|  |  | 100 | 0.009 [0.002, 0.023] | -0.016 [-0.040, 0.000] | 0.024 | - |
|  |  | 1000 | 0.007 [0.001, 0.020] | -0.017 [-0.043, -0.001] | 0.013 | - |
|  | urban areas (R) | 10 | 0.024 [0.004, 0.055] | 0.000 [-0.025, 0.023] | 0.485 | - |
|  |  | 100 | 0.006 [0.000, 0.029] | -0.018 [-0.043, 0.003] | 0.050 | - |
|  |  | 1000 | 0.003 [0.000, 0.023] | -0.021 [-0.045, -0.001] | 0.017 | - |
|  | water areas (C) | 10 | 0.025 [0.011, 0.052] | 0.000 [-0.022, 0.019] | 0.523 | - |
|  |  | 100 | 0.025 [0.011, 0.052] | 0.001 [-0.025, 0.022] | 0.523 | - |
|  |  | 1000 | 0.026 [0.011, 0.055] | 0.002 [-0.025, 0.026] | 0.555 | - |
|  | water areas (R) | 10 | 0.018 [0.003, 0.055] | -0.006 [-0.033, 0.029] | 0.294 | - |
|  |  | 100 | 0.002 [0.000, 0.027] | -0.021 [-0.050, 0.008] | 0.049 | - |
|  |  | 1000 | 0.001 [0.000, 0.025] | -0.022 [-0.050, 0.003] | 0.036 | - |
|  | annual mean | 10 | 0.025 [0.009, 0.049] | -0.001 [-0.012, 0.010] | 0.399 | - |
|  | temperature (C) | 100 | 0.025 [0.009, 0.049] | -0.001 [-0.012, 0.010] | 0.388 | - |
|  |  | 1000 | 0.025 [0.009, 0.049] | -0.001 [-0.012, 0.010] | 0.388 | - |
|  | annual mean | 10 | 0.026 [0.009, 0.053] | 0.001 [-0.010, 0.014] | 0.609 | - |
|  | temperature (R) | 100 | 0.026 [0.009, 0.053] | 0.001 [-0.010, 0.014] | 0.613 | - |
|  |  | 1000 | 0.026 [0.009, 0.053] | 0.001 [-0.011, 0.014] | 0.613 | - |
|  | annual | 10 | 0.025 [0.009, 0.050] | 0.000 [-0.011, 0.012] | 0.452 | - |
|  | precipitation (C) | 100 | 0.025 [0.009, 0.050] | 0.000 [-0.011, 0.012] | 0.455 | - |
|  |  | 1000 | 0.025 [0.009, 0.050] | 0.000 [-0.011, 0.012] | 0.454 | - |
|  | annual | 10 | 0.025 [0.009, 0.051] | 0.000 [-0.019, 0.011] | 0.532 | - |
|  | precipitation (R) | 100 | 0.026 [0.009, 0.051] | 0.000 [-0.010, 0.011] | 0.546 | - |
|  |  | 1000 | 0.026 [0.009, 0.051] | 0.000 [-0.010, 0.011] | 0.546 | - |
| **Circuit-** | elevation (C) | 10 | 0.026 [0.013, 0.052] | -0.018 [-0.029, -0.010] | 0.000 | - |
| **scape** |  | 100 | 0.005 [0.000, 0.026] | -0.038 [-0.062, -0.023] | 0.000 | - |
| **algorithm** |  | 1000 | 0.001 [0.000, 0.015] | -0.041 [-0.074, -0.025] | 0.000 | - |
|  | elevation (R) | 10 | 0.054 [0.033, 0.079] | 0.008 [-0.006, 0.024] | 0.089 | - |
|  |  | 100 | 0.036 [0.019, 0.065] | -0.010 [-0.038, 0.017] | 0.016 | - |
|  |  | 1000 | 0.029 [0.014, 0.057] | -0.015 [-0.045, 0.009] | 0.012 | - |
|  | forest areas (C) | 10 | 0.005 [0.000, 0.020] | -0.038 [-0.064, -0.023] | 0.000 | - |
|  |  | 100 | 0.001 [0.000, 0.010] | -0.042 [-0.075, -0.026] | 0.000 | - |
|  |  | 1000 | 0.001 [0.000, 0.011] | -0.043 [-0.075, -0.025] | 0.000 | - |
|  | forest areas (R) | 10 | 0.033 [0.018, 0.058] | -0.013 [-0.031, 0.008] | 0.006 | - |
|  |  | 100 | 0.027 [0.013, 0.052] | -0.019 [-0.040, 0.006] | 0.004 | - |
|  |  | 1000 | 0.026 [0.012, 0.051] | -0.020 [-0.042, 0.005] | 0.004 | - |
|  | croplands (C) | 10 | 0.048 [0.029, 0.083] | 0.003 [-0.009, 0.011] | 0.070 | - |
|  |  | 100 | 0.035 [0.015, 0.069] | -0.008 [-0.032, 0.012] | 0.021 | - |
|  |  | 1000 | 0.013 [0.002, 0.035] | -0.030 [-0.057, -0.007] | 0.002 | - |
|  | croplands (R) | 10 | 0.029 [0.014, 0.059] | -0.016 [-0.036, -0.001] | 0.002 | - |
|  |  | 100 | 0.001 [0.000, 0.012] | -0.042 [-0.066, -0.024] | 0.000 | - |
|  |  | 1000 | 0.000 [0.000, 0.008] | -0.044 [-0.069, -0.025] | 0.000 | - |
|  | urban areas (C) | 10 | 0.018 [0.008, 0.041] | -0.027 [-0.050, -0.007] | 0.000 | - |
|  |  | 100 | 0.007 [0.002, 0.024] | -0.036 [-0.059, -0.019] | 0.000 | - |
|  |  | 1000 | 0.005 [0.000, 0.019] | -0.039 [-0.065, -0.021] | 0.000 | - |
|  | urban areas (R) | 10 | 0.011 [0.001, 0.031] | -0.034 [-0.059, -0.017] | 0.000 | - |
|  |  | 100 | 0.003 [0.000, 0.014] | -0.040 [-0.073, -0.025] | 0.000 | - |
|  |  | 1000 | 0.002 [0.000, 0.012] | -0.040 [-0.074, -0.026] | 0.000 | - |
|  | water areas (C) | 10 | 0.063 [0.036, 0.095] | 0.014 [-0.020, 0.050] | 0.082 | - |
|  |  | 100 | 0.047 [0.025, 0.081] | 0.000 [-0.035, 0.037] | 0.050 | - |
|  |  | 1000 | 0.042 [0.021, 0.076] | -0.004 [-0.038, 0.031] | 0.042 | - |
|  | water areas (R) | 10 | 0.005 [0.000, 0.020] | -0.039 [-0.072, -0.017] | 0.000 | - |
|  |  | 100 | 0.001 [0.000, 0.009] | -0.042 [-0.075, -0.024] | 0.000 | - |
|  |  | 1000 | 0.001 [0.000, 0.008] | -0.043 [-0.075, -0.024] | 0.000 | - |
|  | annual mean | 10 | 0.042 [0.001, 0.072] | 0.000 [-0.061, 0.018] | 0.048 | - |
|  | temperature (C) | 100 | 0.042 [0.001, 0.071] | 0.000 [-0.061, 0.018] | 0.048 | - |
|  |  | 1000 | 0.042 [0.001, 0.071] | 0.000 [-0.061, 0.018] | 0.048 | - |
|  | annual mean | 10 | 0.044 [0.000, 0.073] | 0.001 [-0.063, 0.020] | 0.052 | - |
|  | temperature (R) | 100 | 0.044 [0.000, 0.073] | 0.001 [-0.063, 0.020] | 0.051 | - |
|  |  | 1000 | 0.044 [0.000, 0.073] | 0.001 [-0.063, 0.020] | 0.051 | - |
|  | annual | 10 | 0.043 [0.000, 0.072] | -0.001 [-0.062, 0.019] | 0.049 | - |
|  | precipitation (C) | 100 | 0.043 [0.000, 0.072] | -0.001 [-0.062, 0.019] | 0.049 | - |
|  |  | 1000 | 0.043 [0.000, 0.072] | -0.001 [-0.062, 0.019] | 0.049 | - |
|  | annual | 10 | 0.044 [0.001, 0.073] | 0.001 [-0.062, 0.021] | 0.051 | - |
|  | precipitation (R) | 100 | 0.044 [0.001, 0.073] | 0.001 [-0.062, 0.021] | 0.051 | - |
|  |  | 1000 | 0.044 [0.001, 0.073] | 0.001 [-0.062, 0.021] | 0.051 | - |
